# Identification of transcription termination defects at DNA hypomethylated transcription termination sites in DNA methyltransferase 3a-deficient vertebrates

**DOI:** 10.1101/2021.06.30.450517

**Authors:** Masaki Shirai, Takuya Nara, Haruko Takahashi, Kazuya Takayama, Yuan Chen, Yudai Hirose, Masashi Fujii, Akinori Awazu, Nobuyoshi Shimoda, Yutaka Kikuchi

## Abstract

CpG methylation in genomic DNA is well known as a repressive epigenetic marker in eukaryotic transcription, and DNA methylation of the promoter regions is correlated with silencing of gene expression. In contrast to the promoter regions, the function of DNA methylation during transcription termination remains to be elucidated. A recent study has revealed that mouse DNA methyltransferase 3a (Dnmt3a) mainly functions in *de novo* methylation in the promoter and gene body regions (including transcription termination sites (TTSs)) during development. To investigate the relationship between DNA methylation overlapping the TTSs and transcription termination, we employed two strategies: informatic analysis using already deposited datasets of *Dnmt3a^-/-^* mouse cells and the zebrafish model system. Bioinformatic analysis using methylome and transcriptome data showed that hypomethylated differentially methylated regions overlapping the TTSs were associated with increased read counts and chimeric transcripts downstream of TTSs in *Dnmt3a^-/-^* Agouti-related protein neurons, but not in *Dnmt3a^-/-^* embryonic stem cells and mouse embryonic fibroblasts. We experimentally detected increased read-through and chimeric transcripts downstream of hypomethylated TTSs in zebrafish maternal-zygotic *dnmt3aa^-/-^* mutant. This study is the first to identify transcription termination defects in DNA hypomethylated TTSs in *Dnmt3a^-/-^* vertebrates.

## Introduction

Transcription by RNA polymerase II (pol II), synthesis process of mRNA from DNA, consists of three biological processes: initiation, elongation, and termination. In higher eukaryotes, two transcription processes (initiation and elongation) and their epigenetic regulation, including DNA methylation and histone modifications, have been extensively studied (Lee and Young 2000; Jones 2012; Gates *et al*. 2017; Chen *et al*. 2018). In transcription termination, it is also well known that most protein-coding genes in eukaryotes possess a polyadenylation signal (PAS) located at the end of the 3’-UTR, and a cleavage and polyadenylation (CPA) complex is assembled around the PAS sequence of nascent pre-mRNA, which directs pol II transcription termination (Proudfoot 2016; Eaton and West 2020). However, in contrast to transcription initiation and elongation, the epigenetic regulation of transcription termination has remained largely unstudied to date.

In eukaryotes ranging from plants to humans, DNA methylation, one of epigenetic regulations, is well characterized as a repressive epigenetic marker and mainly occurs at CpG dinucleotides (Li and Zhang 2014; SCHÜBELER 2015; Luo *et al*. 2018). Advances in sequencing technology have resulted in genomic methylation profiling at single-base resolution revealing that the global DNA methylation (5-methylcytosine, 5mC) is seen in mammals, except for short CpG-rich regions known as CpG islands (Li and Zhang 2014; SCHÜBELER 2015; Luo *et al*. 2018). Studies of methylome profiling have shown that DNA methylation patterns dynamically change during development and disease and that these patterns are linked to gene expression in mammals (Smith and Meissner 2013; Greenberg and Bourc’his 2019). Genome-wide methylome analysis revealed that the DNA methylation levels around the promoter regions, including the transcription start site (TSS), are negatively correlated with gene expression (Ball *et al*. 2009; Rauch *et al*. 2009). Conversely, DNA methylation levels in the gene bodies are positively correlated with gene expression (Kulis *et al*. 2013; Teissandier and Bourc’his 2017). Although two possible mechanisms, regulation of splicing and prevention of cryptic promoters, have been proposed for methylation function in the gene body (Kulis *et al*. 2013; Teissandier and Bourc’his 2017), the significance of DNA methylation around transcription termination sites (TTSs; cleavage/polyadenylation sites) remains unresolved.

The DNA methylation patterns are well known to be established by the DNA methyltransferases (Dnmts), which are categorized into two types in mammals: maintenance Dnmt (Dnmt1) and *de novo* Dnmts (Dnmt3a and Dnmt3b) (Li *et al*. 1992; Okano *et al*. 1999; Li and Zhang 2014). Knock-out mice of both Dnmt3a and Dnmt3b, which die at embryonic day 9.5, fail to start *de novo* methylation after implantation (Okano *et al*. 1999), and genetic mutation of Dnmt3a or Dnmt3b leads to various human diseases (Greenberg and Bourc’his 2019), suggesting the critical role of DNA methylation in development and homeostasis in mammals through the regulation of gene expression. A very recent study using mouse embryonic fibroblasts (MEFs) with genetic ablation of *de novo* Dnmts reported that Dnmt3a functions in *de novo* methylation at both TSS regions (TSS ± 1000 bp) and gene bodies (from TSS regions to TTS) of the Polycomb group (PcG) target developmental genes, whereas Dnmt3b mainly functions in X-chromosome methylation (Yagi *et al*. 2020). These results indicate that although the enzymatic functions of both proteins are redundant, genomic target regions are distinct in mammals. Although mammalian Dnmt3a mutant phenotypes and target regions have been reported, the role of methylation during transcription termination by Dnmt3a has not yet been reported.

In the present study, we used bioinformatic analysis and a zebrafish model animal system to explore the function of Dnmt3a-mediated hypomethylation at the TTSs. The bioinformatic analysis showed that hypomethylated differentially methylated regions (hypoDMRs) overlapping the TTSs were associated with increased read counts and chimeric transcripts downstream of TTSs in *Dnmt3a^-/-^* Agouti-related protein neurons, but not in *Dnmt3a^-/-^* embryonic stem cells (ESCs) and mouse embryonic fibroblasts (MEFs). Based on bioinformatic analysis, to experimentally confirm the transcription termination defects in *Dnmt3a^-/-^* vertebrates, we generated zebrafish maternal-zygotic *dnmt3aa^-/-^* (MZ*dnmt3aa^-/-^*) mutants and identified an increase in both read-through and chimeric transcripts in the hypoDMRs overlapping the TTSs in MZ*dnmt3aa^-/-^* mutant. Taken together, in *Dnmt3a*-deficient vertebrates, we identified transcription termination defects in DNA hypomethylated TTSs.

## Materials and methods

### Zebrafish experiments, ethic statement, and generation of MZ*dnmt3aa^-/-^* mutant fish using CRISPR/Cas9 system

Adult zebrafish and zebrafish larvae were maintained as previously described (Westerfield 1993). All zebrafish experiments were approved from the Hiroshima University Animal Research Committee (Permit Number: F18-2-7).

Genome editing using the CRISPR/Cas9 system was performed as previously described (Ansai and Kinoshita 2014). Briefly, the pDR274 vector (Addgene Plasmid 42250) was digested with *Bsa*I, and then annealed oligonucleotides for sgRNAs were cloned into the pDR274 vector. The sequences of the oligonucleotides for sgRNAs are listed in Supplementary Table S1. The sgRNA expression vectors were digested by *Dra*I, and the sgRNAs were synthesized using the MAXIscript^TM^ T7 Kit (Invitrogen, Thermo Fisher Scientific). The synthesized sgRNAs were purified using the mirVana^TM^ miRNA Isolation Kit (Invitrogen, Thermo Fisher Scientific). In addition, the Cas9 expression vector (Addgene Plasmid 51815) was linearized by *Not*I and *Cas9* mRNA was synthesized using the mMESSAGE mMACHINE^TM^ SP6 Kit (Invitrogen, Thermo Fisher Scientific). A mixture of three types of sgRNAs for the zebrafish *dnmt3aa* gene and *Cas9* mRNA of appropriate concentrations was injected at the one-cell stage. To identify the mutations, the target site of the CRISPR/Cas9 system on *dnmt3aa* gene was amplified using TaKaRa Ex Taq^®^ (Takara Bio) with the primers listed in Supplementary Table S2. The amplicons were purified using the QIAquick PCR Purification Kit (QIAGEN) and then sequenced using the BigDye^TM^ Terminator v3.1 Cycle Sequencing Kit (Applied Biosystems, Thermo Fisher Scientific) and primers listed in Supplementary Table S2.

The amputated fins of *dnmt3aa^-/-^* mutant were digested overnight in lysis buffer (100 mM NaCl, 20 mM Tris-HCl pH 8, 50 mM EDTA pH 8, 0.5% SDS and 36 ug/ml Proteinase K) at 55°C. Proteinase K was inactivated at 98°C for 10 min. A heteroduplex mobility assay (HMA) was performed to detect the *dnmt3aa* mutation. The target site of the CRISPR/Cas9 system on exon 1 of the *dnmt3aa* gene was amplified using BIOTAQ^TM^ DNA Polymerase (Meridian Bioscience); the primers used are listed in Supplementary Table S2. Amplicons were analyzed using polyacrylamide gel electrophoresis. MZ*dnmt3aa^-/-^* mutants were obtained by crossing *dnmt3aa^-/-^* adults.

### Whole-genome bisulfite library preparation and sequencing of zebrafish samples

Zebrafish wild-type (WT) and MZ*dnmt3aa^-/-^* mutant embryos at 2 days post-fertilization (dpf) were digested with proteinase K and sodium dodecyl sulfate to extract genomic DNA. DNA samples were prepared by mixing zebrafish genomic DNA and unmethylated lambda DNA (Promega) as spike control, at a ratio of 1000:1. DNA samples (100 ng) were subjected to bisulfite treatment using the EZ DNA Methylation-Gold Kit (Zymo Research). Bisulfite-treated DNA samples (50 ng) were subjected to 10 cycles of PCR using random primers from the TruSeq DNA Methylation Kit (Illumina) to add adapters. Libraries with adapters were sequenced on the HiSeq X Five Sequencing System (Illumina) with 150 bp single-end reads. Bisulfite treatment, library preparation, and sequencing were performed by Takara Bio Inc. (Shiga, Japan).

### Whole-genome bisulfite sequencing (WGBS) data analysis of mouse dataset

The publicly available datasets used in this study were from the Gene Expression Omnibus (GEO), and the accession numbers are GSE100957 (J1 mESCs) (Gu *et al*. 2018), GSE111172 (MEFs) (Yagi *et al*. 2020), and GSE122405 (AgRPs) (Mackay *et al*. 2019). The FASTQ-format read sequences were trimmed of adaptors and low-quality bases were filtered using Trim Galore (version 0.6.4) (https://github.com/FelixKrueger/TrimGalore) with Cutadapt (version 2.10) (Martin 2011) and FastQC (version 0.11.8) (https://www.bioinformatics.babraham.ac.uk/projects/fastqc/). After trimming, the reads were aligned to the mouse genome (mm10) using Bismark (version 0.20.0) (Krueger and Andrews 2011) and Bowtie (version 1.0.0) (Langmead *et al*. 2009). The aligned reads were followed by the removal of PCR duplicates using deduplicate_bismark (version 0.20.0) (Krueger and Andrews 2011), and methylation calls were performed using bismark_methylation_extractor (version 0.20.0) (Krueger and Andrews 2011). The reads from samples with multiple sequencing lanes were merged using Samtools (version 1.9) (Li *et al*. 2009) after removing the PCR duplicates. The methylation ratio of CpG sites with a coverage of over 5 reads in WT and *Dnmt3a^-/-^* was calculated from the methylation call data. The methylation ratio was visualized using Integrative Genomics Viewer (IGV) (version 2.7.2) (Robinson *et al*. 2011).

### WGBS data analysis of zebrafish datasets

The read sequences in FASTQ format were first trimmed by removing six bases from random primers of TruSeq DNA Methylation Kit (Illumina) and the last 50 bases of low-quality reads. After trimming, the reads were aligned to the zebrafish genome (danRer10) and lambda phage genome using Bismark (version 0.10.1) (Krueger and Andrews 2011) and Bowtie (version 1.0.0) (Langmead *et al*. 2009). Once aligned, reads corresponding to PCR duplicates were removed and methylation calls were performed. The bisulfite conversion rates based on spike control for all samples were over 99%, and the error rates were as follows: WT, 0.3% and MZ*dnmt3aa^-/-^* mutant, 0.2%. Statistically significant methylated cytosine sites were identified using a binomial distribution with a false discovery rate ≤ 0.05, calculated by the Benjamini-Hochberg (BH) method, and read depth ≥ 5. Data analysis was performed by Takara Bio, Inc. (Shiga, Japan) as previously described (Lister *et al*. 2009). WGBS data were deposited in the GEO under accession number GSE178691. The methylation ratio of CpG sites with a coverage of over 5 reads in WT and MZ*dnmt3aa^-/-^* mutant as calculated from the methylation call data. The methylation ratio was visualized using the IGV (version 2.7.2) (Robinson *et al*. 2011).

### Identification of differentially methylated regions (DMRs)

The methylation call data of mice and zebrafish were used to identify DMRs using swDMR (version 1.0.0) (Wang *et al*. 2015) with the followed settings: cytosine type, CG; window, 500; step size, 50; points (lowest number of selected cytosine type in the window), 3; coverage, 5; fold (methylation level difference), 1.5; diff (value of max-min methylation level), 0.1; p-value, 0.05; fdr, 0.05. Samples were compared using the Fisher’s test. DMRs overlapped genomic positions (TSS, gene body, TTS, and intergenic region) were determined using bedtools (version 2.28.0) (Quinlan and Hall 2010) intersect. The DMRs that overlap both TSS and TTS were included in the DMRs that overlap TTS. Heatmaps, violin plots, scatterplots with marginal distribution, and horizontal bar graphs were generated from swDMR output using Python and R scripts.

### RNA sequencing (RNA-seq) library preparation and sequencing of zebrafish samples

Tissue lysates were prepared from 30 pooled 2 dpf WT and MZ*dnmt3aa^-/-^* mutant by homogenization using TissueLyser LT (QIAGEN) with zirconia beads for 10 s at 50 Hz. RNA was extracted from the lysate using the PureLink^®^ RNA Mini Kit (Thermo Fisher Scientific). RNA quality of all samples was RNA Integrity Number (RIN) > 9.0, using the 4150 TapeStation System (Agilent). Total RNA was treated using the NEBNext^®^ Poly(A) mRNA Magnetic Isolation Module (New England Biolabs) for mRNA isolation. Strand-specific RNA-seq libraries were constructed using the NEBNext^®^ Ultra^TM^ Directional RNA Library Prep Kit for Illumina^®^ (New England Biolabs). The libraries with adapters were sequenced on the Novaseq 6000 (Illumina) with 2 × 150 bp paired-end reads. RNA extraction, mRNA isolation, library preparation, and sequencing were performed by Rhelixa Inc. (Tokyo, Japan).

### RNA-seq data analysis for mouse and zebrafish datasets

The publicly available datasets of mice deposited in the GEO and the generated datasets of zebrafish were used in this study. Accession numbers are GSE100957 (J1 mESCs) (Gu *et al*. 2018), GSE84165 (MEFs) (Yagi *et al*. 2017a; Yagi *et al*. 2019), GSE111172 (MEFs) (Yagi *et al*. 2020), and GSE122405 (AgRPs) (Mackay *et al*. 2019). RNA-seq data of zebrafish were deposited in the GEO under accession number GSE178691. FASTQ files from samples with multiple sequencing lanes were merged. The FASTQ-format sequence reads were trimmed of adaptors and low-quality bases were filtered using Trim Galore (version 0.6.4) (https://github.com/FelixKrueger/TrimGalore) with Cutadapt (version 1.18 or 2.10) (Martin 2011) and FastQC (version 0.11.8 or 0.11.9) (https://www.bioinformatics.babraham.ac.uk/projects/fastqc/). After trimming, the reads were aligned to the mouse genome (mm10) and the zebrafish genome (danRer10) using HISAT2 (version 2.1.0) (Kim *et al*. 2015). Alignment of zebrafish datasets was performed using the strand-specific option. Aligned files (.bam) were sorted and indexed using Samtools (version 1.9) (Li *et al*. 2009). In addition, the strand-specific aligned files of zebrafish were generated using Samtools (version 1.9) (Li *et al*. 2009). Read coverage was normalized with 1 bin size by counts per million mapped reads (CPM) using the bamCoverage tool from the DeepTools (version 3.3.2) (RAMÍREZ *et al*. 2016) and visualized using the IGV (version 2.7.2) (Robinson *et al*. 2011). Splice junction data of chimeric transcripts in the aligned files were inspected manually using the IGV (version 2.7.2) (Robinson *et al*. 2011).

### Calculation of expression level downstream of TTS from RNA-seq data

Quantification of expression levels was completed using StringTie (version 2.1.2) (Pertea *et al*. 2015) with NCBI RefSeq annotation. The expression values of each transcript were normalized to the transcripts per million (TPM). Transcripts expressed in WT and *Dnmt3a^-/-^* mouse cells or MZ*dnmt3aa^-/-^* mutant were extracted from all transcripts (TPM > 0). In addition, the transcripts expressed in both strains and overlapped hypoDMRs across the TTS were extracted.

All reads that overlapped the exonic or intronic sequences in the aligned files (.bam) were removed using bedtools (version 2.28.0) (Quinlan and Hall 2010) intersect with the NCBI RefSeq annotation. In addition, the intergenic reads were counted within 3 kb downstream of the TTS using bedtools (version 2.28.0) (Quinlan and Hall 2010) multicov, except when the genomic region downstream of the TTS was less than 3 kb. The expression level was calculated using the following equation (TPM 3 kb):

TPM 3 kb = Number of reads aligned in 3kb downstream of TTS of each transcript × 10^6^ / Total number of reads aligned in 3kb downstream of TTS.

Normalized read counts were compared in WT and *Dnmt3a^-/-^* mouse cells or MZ*dnmt3aa^-/-^* zebrafish mutants as a metagene analysis.

Box plots were generated using R scripts, and statistical significance was determined using Brunner-Munzel test (*p* ≤ 0.05).

### Calculation of expression level of chimeric transcripts from RNA-seq data

Chimeric transcripts, which were aberrantly fused with exons of downstream genes or intergenic DNA, were manually counted using IGV (Robinson *et al*. 2011). The expression level was calculated as the counts per million (CPM) mapped reads. Normalized read counts were compared in WT and *Dnmt3a^-/-^* mouse cells or MZ*dnmt3aa^-/-^* zebrafish mutant as a metagene analysis. Transcripts whose expression increased downstream of the TTS were used for metagene analysis. Box plots were generated using R scripts. Statistical significance was determined using the Brunner-Munzel test (*p* ≤ 0.05).

### RNA preparation, quantitative PCR (qPCR), and sequencing for identification of read-through and chimeric transcripts of zebrafish samples

Total RNA was extracted using TRIzol (Invitrogen, Thermo Fisher Scientific) from six samples of 30 pooled 2 dpf WT and MZ*dnmt3aa^-/-^* mutant, and the six extracted total RNA samples were combined as one sample. Total RNA (150 ug) was treated using the *Oligotex*^TM^ -*dT30*<*Super*> mRNA Purification Kit (Takara Bio). The purified poly(A)^+^ RNA was reverse-transcribed with oligo(dT) primer and reverse transcriptase XL (AMV) (Takara Bio).

qPCR was performed using the StepOne^TM^ Real-Time PCR System and PowerUp^TM^ SYBR^TM^ Green Master Mix (Applied Biosystems, Thermo Fisher Scientific). Relative expression levels were calculated by the ΔΔCT method, and targets were normalized by the mean value of a control gene, *ubiquitin-conjugating enzyme E2A* (*ube2a*). The qPCR primers used are shown in Supplementary Table S3. Bar charts were generated from the mean values and standard error (SE) data using R scripts. Statistical significance was determined using the t-test (*p* ≤ 0.05).

Sequencing of chimeric transcript was performed using MZ*dnmt3aa^-/-^* mutant. The product of *im:7152937*-intergenic DNA (iD)-*meiosis-specific nuclear structural 1* (*mns1*) chimeric transcript was amplified using BIOTAQ^TM^ DNA Polymerase (Meridian Bioscience) with the primers listed in Supplementary Table S4. The amplicons were purified using the QIAquick Gel Extraction Kit (QIAGEN) after agarose gel electrophoresis. The purified product was subcloned using a TA-cloning Kit (Thermo Fisher Scientific), and the plasmid DNA was sequenced using the BigDye^TM^ Terminator v3.1 Cycle Sequencing Kit (Applied Biosystems, Thermo Fisher Scientific) and primers listed in Supplementary Table S4.

## Results

### Genomic characteristics of hypoDMRs overlapping the TTS in *Dnmt3a*-deficient mouse cells

To investigate the functional relevance between DNA methylation and transcription termination, we searched datasets of *Dnmt3a*-deficient mammalian cells that contain both methylome (WGBS) and transcriptome (RNA-seq) data. Five datasets of WGBS (human: one, mouse: four) and ten datasets of RNA-seq (human: four, mouse: six) were obtained from GEO. Four datasets (human: one, mouse: three) contained both WGBS and RNA-seq, and among them we chose three types of mouse datasets (J1 mESCs (Gu *et al*. 2018); MEFs (Yagi *et al*. 2020); and Agouti-related protein neurons, AgRPs (Mackay *et al*. 2019)) because of the differences in their cell differentiation states. The outline of bioinformatics analysis and experimental design was shown in Supplementary Figure S1. We extracted the hypermethylated DMRs (hyperDMRs) and hypoDMRs from these datasets and found that the number of hypoDMRs in *Dnmt3a*^-/-^ J1 mESCs were approximately 57 and 181 times greater than that of *Dnmt3a*^-/-^ MEFs and AgRPs, respectively (Supplementary Figure S2, A-C). The hypoDMRs were categorized by genomic position (gene body, intergenic region, and gene body & intergenic region) (Supplementary Figure S3). The number of hypoDMRs gradually decreased in the order of *Dnmt3a*^-/-^ J1 mESCs, MEFs, and AgRPs, whereas their percentages of genomic positions were nearly the same among these *Dnmt3a*-deficient cells (Supplementary Figure S3). Next, we selected the hypoDMRs overlapping TTSs in three types of *Dnmt3a*^-/-^ mouse cells (Figure 1, A and B) and found that the number of hypoDMRs overlapping the TTSs in *Dnmt3a*^-/-^ MEFs or AgRPs was less than 1% or 0.4% of that of *Dnmt3a*^-/-^ J1 mESCs, respectively (Figure 1C). These data suggest that DNA methylation regulated by Dnmt3a may be correlated with the differentiation state of mouse cells. Two-dimensional (2D) scatterplots with marginal distribution showed that the hypoDMRs overlapping the TTSs were almost symmetrically distributed across the TTSs in the *Dnmt3a*^-/-^ J1 mESCs and MEFs, whereas the distribution of hypoDMRs was slightly shifted upstream of the TTSs in *Dnmt3a*^-/-^ AgRPs (Figure 1D).

**Figure 1.**
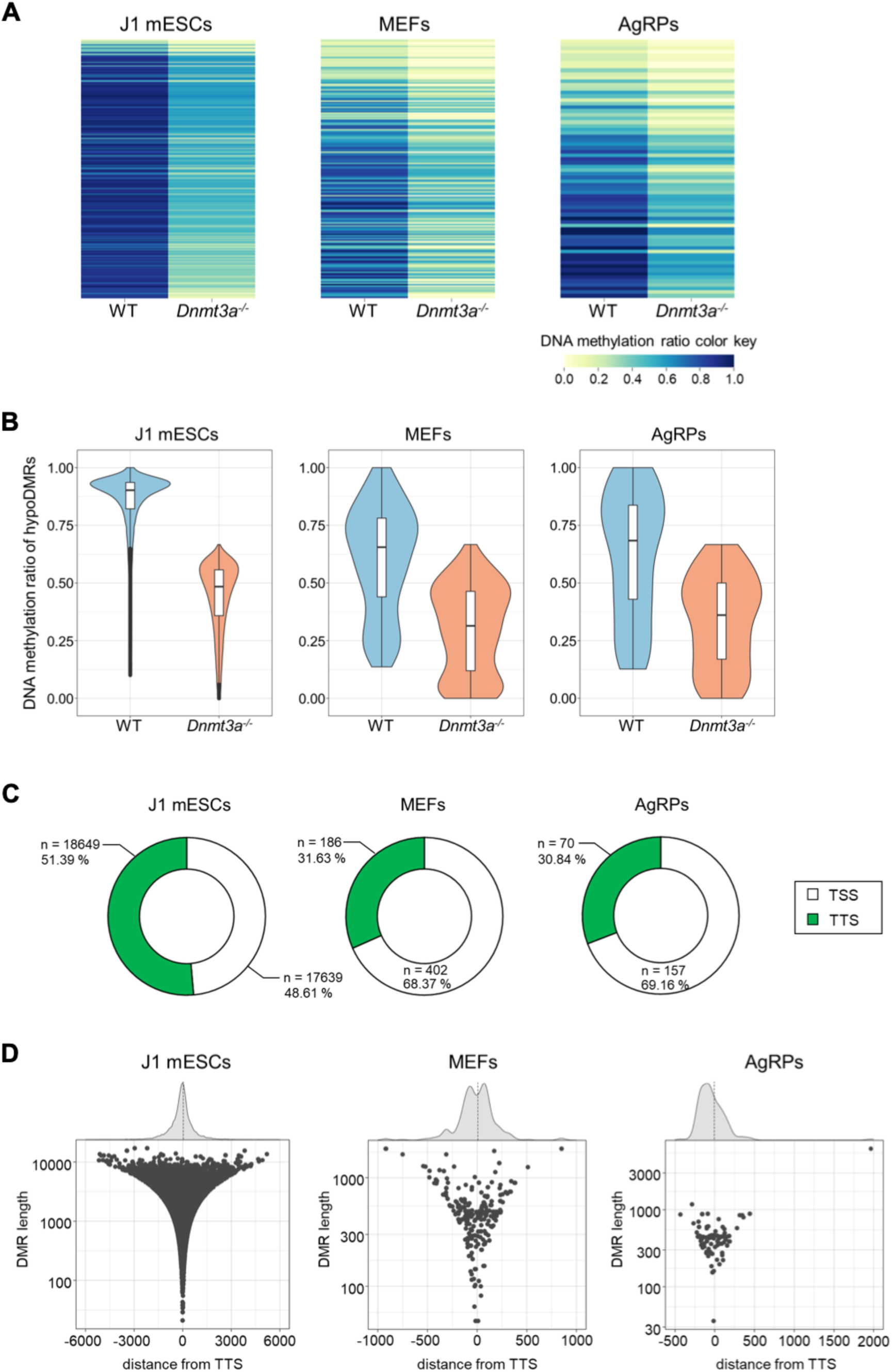
Characterization of hypoDMRs in three *Dnmt3a*-deficient mouse cells. (A) Heatmaps of DNA methylation profiles of hypoDMRs overlapping TTSs comparing WT with *Dnmt3a^-/-^* mutants (J1 mESCs, MEFs, and AgRP). Color scale indicates DNA methylation ratio. (B) Violin plots showing DNA methylation ratio of the hypoDMRs overlapping the TTSs comparing WT with *Dnmt3a^-/-^* mutants (J1 mESCs, MEFs, and AgRP). (C) Pie charts depicting the number and percentage of hypoDMRs overlapping the TSSs and TTSs in *Dnmt3a^-/-^* J1 mESCs, MEFs, and AgRPs. (D) 2D scatterplots with marginal distribution comparing hypoDMR length versus distance from TTS in *Dnmt3a^-/-^* J1 mESCs, MEFs, and AgRPs. Dots represent the center of hypoDMRs.

### HypoDMRs overlapping the TTSs are associated with an increased read counts and chimeric transcripts downstream of TTSs in *Dnmt3a^-/-^* AgRPs

To explore the effect of hypomethylation overlapping the TTSs on transcription termination, we analyzed RNA-seq data from three different types of *Dnmt3a*^-/-^ mouse cells. Read counts aligned in 3 kbp downstream of TTSs were normalized by TPM (3 kb) in wild-type (WT) and *Dnmt3a*^-/-^ J1 mESCs, MEFs, and AgRPs. Metagene analysis showed that the read counts downstream of TTSs increased significantly in *Dnmt3a*^-/-^ AgRPs compared to those in WT cells (Figure 2, A-C and Supplementary Figure S4A; *Ring Finger 157* (*Rnf157*) gene as an example of increased read counts), and the higher expression downstream of TTSs was observed in more than half of the genes with hypoDMRs in mutant cells (Figure 2D). The centers of the hypoDMRs which cause an increase in read counts downstream of TTSs were distributed close to the TTSs in *Dnmt3a*^-/-^ AgRPs (Supplementary Figure S5, A, A’, and B). Conversely, the read counts in *Dnmt3a*^-/-^ J1 mESCs and MEFs were unchanged compared to those in WT cells (Figure 2, A and B).

**Figure 2.**
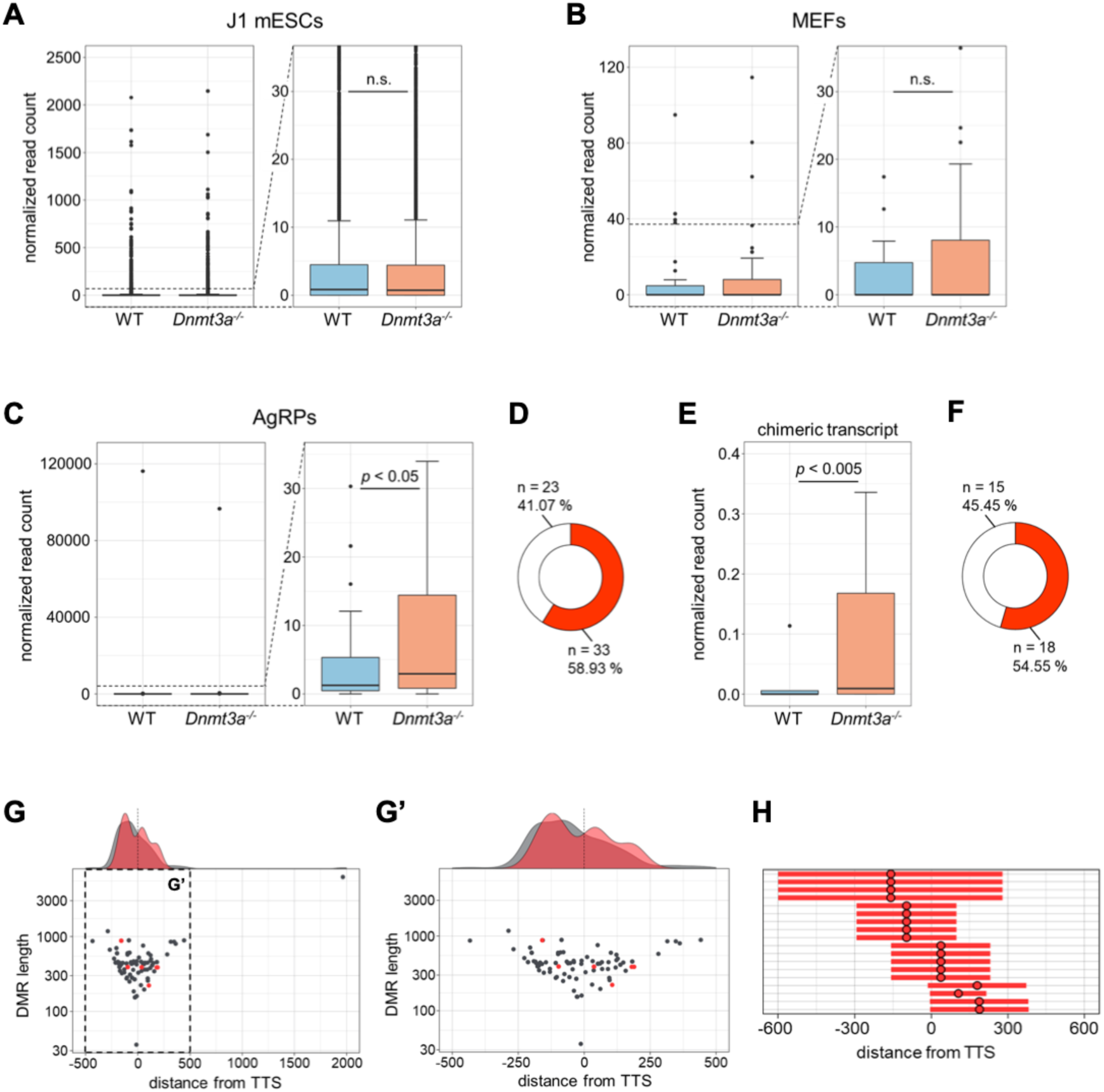
Metagene analysis of transcripts downstream of TTSs in three *Dnmt3a*-deficient mouse cells. (A-C) Box plots showing the normalized read counts in WT and *Dnmt3a^-/-^* mutants (J1 mESCs, MEFs, and AgRPs). Right graphs are enlarged views of the left ones, respectively. (D) A pie chart depicting the number and percentage of transcripts with increased read counts (red) or decreased/unchanged read counts (white) in *Dnmt3a^-/-^* AgRPs. (E) A box plot showed chimeric transcripts in WT and *Dnmt3a^-/-^* AgRPs. (F) A pie chart depicting the number and percentage of transcripts with increased chimeric transcripts (red) or decreased/unchanged chimeric transcripts (white) in *Dnmt3a^-/-^* AgRPs. (G, G’) 2D scatterplots with marginal distribution comparing hypoDMR length versus distance from TTS in *Dnmt3a^-/-^* AgRPs. Grey and red dots represent the center of hypoDMRs. Red dots represent hypoDMRs, which lead to increased chimeric transcripts. A right graph (G’) is an enlarged view of the left one (G). (H) A horizontal bar graph represents the distance from TTS of hypoDMRs, which lead to increased chimeric transcripts, relative to TTS. Red dots and bars represent the center and range of hypoDMRs relative to TTS position, respectively.

We then analyzed the content of transcripts downstream of hypomethylated TTSs in the *Dnmt3a*^-/-^ AgRPs, and found chimeric transcripts that were aberrantly fused with exons of downstream genes or intergenic DNA (iD) in *Dnmt3a*^-/-^ AgRPs, as observed in WT cells. Frequency of chimeric transcripts (about 54.6%) increased significantly in the *Dnmt3a*^-/-^ AgRPs compared to those in WT cells (Figure 2, E and F). We showed that splicing of the *Rnf157* as an example of chimeric transcripts in the *Dnmt3a*^-/-^ AgRPs, and found aberrant transcripts that connect an *Rnf157* exon either to the intergenic DNA or to *Unkempt family zinc finger* (*Unk*) gene (Supplementary Figure S4B). The centers of DMRs causing these chimeric transcripts were distributed close to the TTS (Figure 2, G, G’, and H). In summary, the increase in the chimeric transcripts in the *Dnmt3a*^-/-^ AgRPs clearly shows that transcription termination is defective and that read-through occurs at hypomethylated TTSs.

### Genomic characteristics of hypoDMRs overlapping the TTSs in MZ*dnmt3aa^-/-^* mutant

To explore the correlation between the transcription termination defects and DNA methylation in an animal model, we examined the methylome and transcriptome analysis using the zebrafish as a model. Two zebrafish orthologs (*dnmt3aa* and *dnmt3ab*) of mammalian Dnmt3a have been previously identified (Takayama *et al*. 2014), and the detailed expression profiles of *dnmt3aa* during development and regeneration showed that *dnmt3aa* was maternally expressed and its expression was observed in more organs than *dnmt3ab* (Takayama *et al*. 2014). Therefore, we attempted to generate the MZ*dnmt3aa^-/-^* knockout mutant by using CRISPR/Cas9 method (Figure 3A) and found that they were viable, fertile and showed no specific phenotype at 2 dpf (Figure 3B). Analyses of WGBS and RNA-seq were performed at 2 dpf, because most of the organs had already formed at this stage. In MZ*dnmt3aa^-/-^* mutant, 43.6% DMRs were hypomethylated (Supplementary Figure S6) and the hypoDMRs were categorized by genomic position (gene body, intergenic region, and gene body & intergenic region) (Supplementary Figure S7). We then selected the hypoDMRs overlapping the TTSs, and 27 hypoDMRs were identified overlapping the TTSs (Figure 3, C-E). A 2D scatterplot with marginal distribution showed that the hypoDMRs overlapping the TTSs were almost symmetrically distributed across the TTSs in MZ*dnmt3aa^-/-^* mutant (Figure 3, F and F’), as observed in *Dnmt3a*^-/-^ J1 mESCs and MEFs.

**Figure 3.**
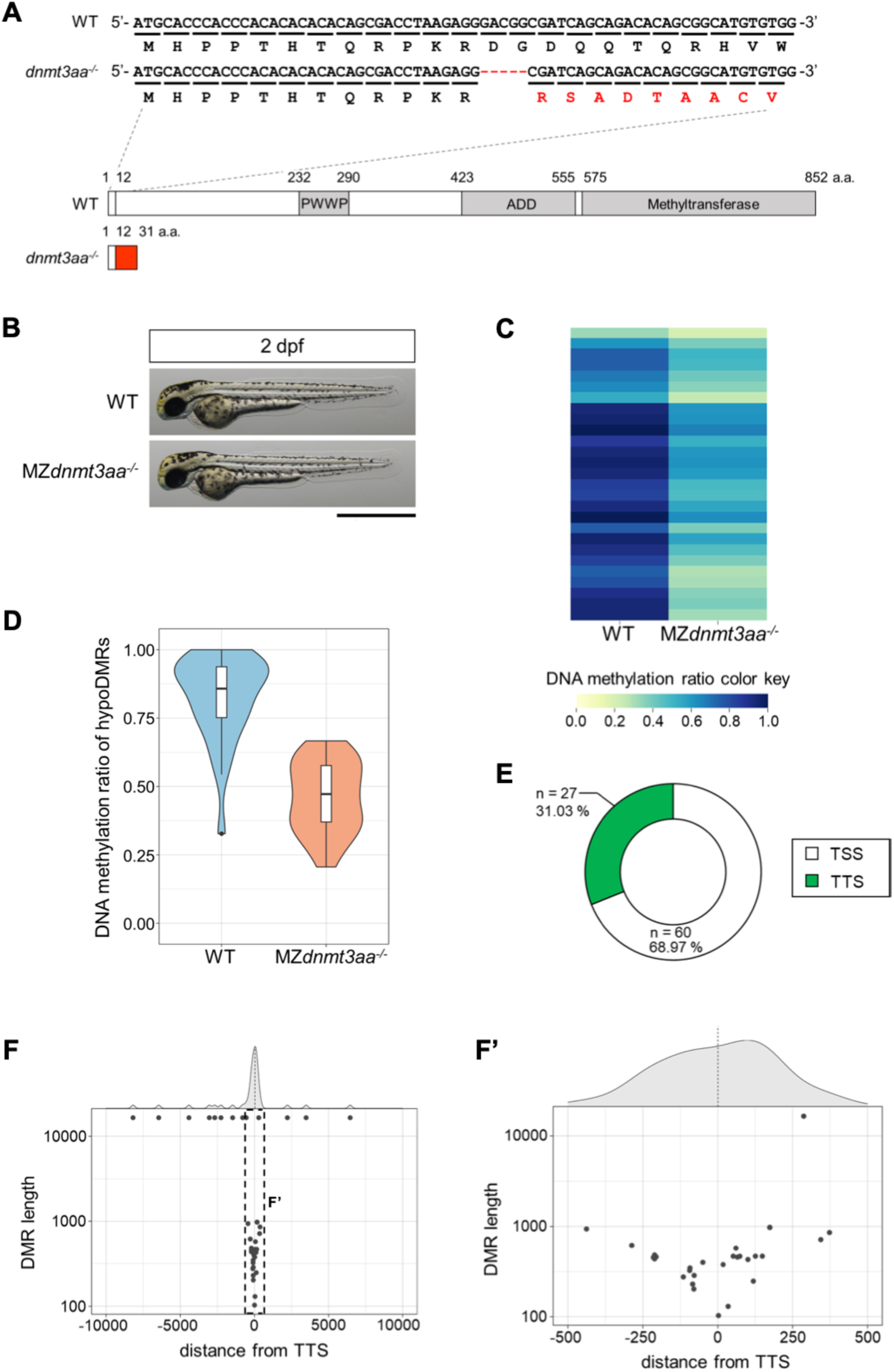
Generation of MZ*dnmt3aa^-/^*^-^ zebrafish mutant and characterization of hypoDMRs. (A) The *dnmt3aa* genomic DNA sequence and CRISPR/Cas9-mediated indel mutation, with predicted amino acid sequences, in WT and MZ*dnmt3aa^-/-^* mutant are shown. The MZ*dnmt3aa^-/-^* mutant has a 5-bp deletion which leads to a frame-shift at amino acid 13 position. Schematic diagram shows predicted protein products from the WT and the mutant allele. It is speculated that the deletion mutation results in the production of a truncated protein with additional sequences (red box) at the C-termini. Putative functional domains (PWWP, cysteine-rich ATRX-Dnmt3-Dnmt3L (ADD), and methyltransferase domains) in the WT protein are represented as grey colored boxes. (B) Gross morphology of the MZ*dnmt3aa^-/-^* mutant is identical to WT at 2 dpf. Scale bar: 1 mm. (C) A heatmap of DNA methylation profiles of hypoDMRs overlapping TTSs comparing WT with MZ*dnmt3aa^-/-^* mutant. Color scale indicates DNA methylation ratio. (D) A violin plot showing DNA methylation ratio of the hypoDMRs overlapping the TTSs comparing WT with MZ*dnmt3aa^-/-^* mutant. (E) A pie chart depicting the number and percentage of hypoDMRs overlapping the TSSs and TTSs in MZ*dnmt3aa^-/-^* mutant. (F, F’) 2D scatterplots with marginal distribution comparing hypoDMR length versus distance from TTS in MZ*dnmt3aa^-/-^* mutant. The right graph (F’) is an enlarged view of the left one (F). Grey dots represent the center of hypoDMRs.

### Increases in both chimeric and read-through transcripts were identified in the hypoDMRs overlapping TTSs in MZ*dnmt3aa^-/-^* mutant

We also analyzed the RNA-seq data of MZ*dnmt3aa*^-/-^ mutant and the read counts from genes containing hypoDMRs overlapping the TTSs, which were normalized in the WT and mutant. Metagene analysis showed that the read counts downstream of the TTSs in MZ*dnmt3aa*^-/-^ mutant were unchanged compared to those in WT (Figure 4A). The transcripts that showed an increase (30.3%) were almost symmetrically distributed across the TTSs in the mutant (Figure 4B and Supplementary Figure S8). Next, we have extracted chimeric transcripts fused with exons of downstream genes and/or intergenic DNA (iD). Normalized read counts of the chimeric transcripts (30%) were also unchanged in MZ*dnmt3aa*^-/-^ mutant compared to those in WT (Figure 4, C and D). Three increased chimeric transcripts (*cytokine like 1* (*cytl1*), one; *im:7152937*, two) were found in two different DMRs, which were close to TTSs (Figure 4, E, E’, and F and Supplementary Figure S9 and S10).

**Figure 4.**
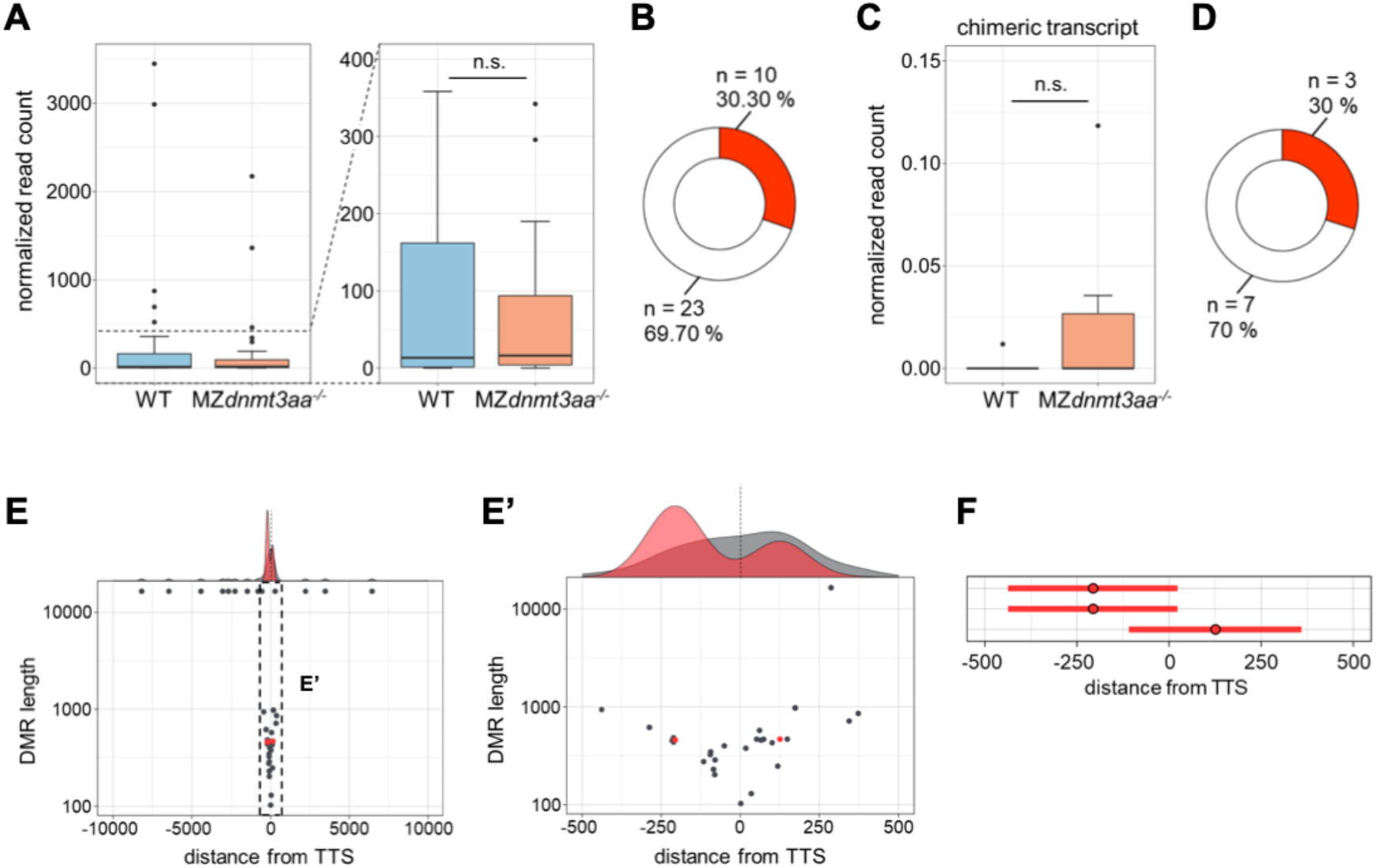
Metagene analysis of transcripts downstream of TTSs in MZ*dnmt3aa^-/-^* mutant. (A) A box plot showing the normalized read counts in WT and MZ*dnmt3aa^-/-^* mutant. The right graph is an enlarged view of the left one. (B) A pie chart depicting the number and percentage of transcripts with increased read counts (red) or decreased/unchanged read counts (white) in MZ*dnmt3aa^-/-^* mutant. (C) A box plot showing chimeric transcripts in WT and MZ*dnmt3aa^-/-^* mutant. (D) A pie chart depicting the number and percentage of transcripts with increased chimeric transcripts (red) or decreased/unchanged chimeric transcripts (white) in MZ*dnmt3aa^-/-^* mutant. (E, E’) 2D scatterplots with marginal distribution comparing hypoDMR length versus distance from TTS in MZ*dnmt3aa^-/-^* mutant. Grey and red dots represent the center of hypoDMRs. Red dots represent hypoDMRs, which lead to increased chimeric transcripts. The right graph (E’) is an enlarged view of the left one (E). (F) A horizontal bar graph represents the distance from TTS of hypoDMRs, which lead to increased chimeric transcripts, relative to TTS. Red dots and bars represent the center and range of hypoDMRs relative to TTS position, respectively.

To further analyze the transcription termination defect caused by hypoDMRs overlapping the TTSs, we examined the detailed expression downstream of and across the TTSs of three genes that had shown high expression levels downstream of TTSs (*cytl1*, *im:7152937*, and *potassium channel, subfamily K, member 1b* (*kcnk1b*)) by RNA-seq. The expression level of three transcripts (*cytl1*_doT, *im:7152937*_doT, and *kcnk1b*_doT) in MZ*dnmt3aa^-/-^* mutant had significantly increased downstream of the TTSs compared to that in WT, as detected by qPCR (Figure 5, A and E). We also found a significant increase in the two transcripts (*im:7152937*_acT and *kcnk1b*_acT) across the TTSs (Figure 5, B, F, and G), demonstrating that read-through occurs in these genes. In addition to read-through, chimeric transcripts (*im:7152937*-iD_cht and *im:7152937*-iD-*meiosis-specific nuclear structural 1* (*mns1*)_cht) were detected in *im:7152937* gene (Supplementary Figure S10 and S11), and qPCR analysis showed that the expression level of chimeric transcripts between *im:7152937* and iD or *mns1* had increased significantly in MZ*dnmt3aa^-/-^* mutant (Figure 5C, D, and H). Taken together, the read-through and chimeric transcripts of three genes in MZ*dnmt3aa^-/-^* mutant, combined with the results from the *Dnmt3a*^-/-^ AgRPs analyses, clearly support our idea that transcription termination is defective in hypomethylated TTSs.

**Figure 5.**
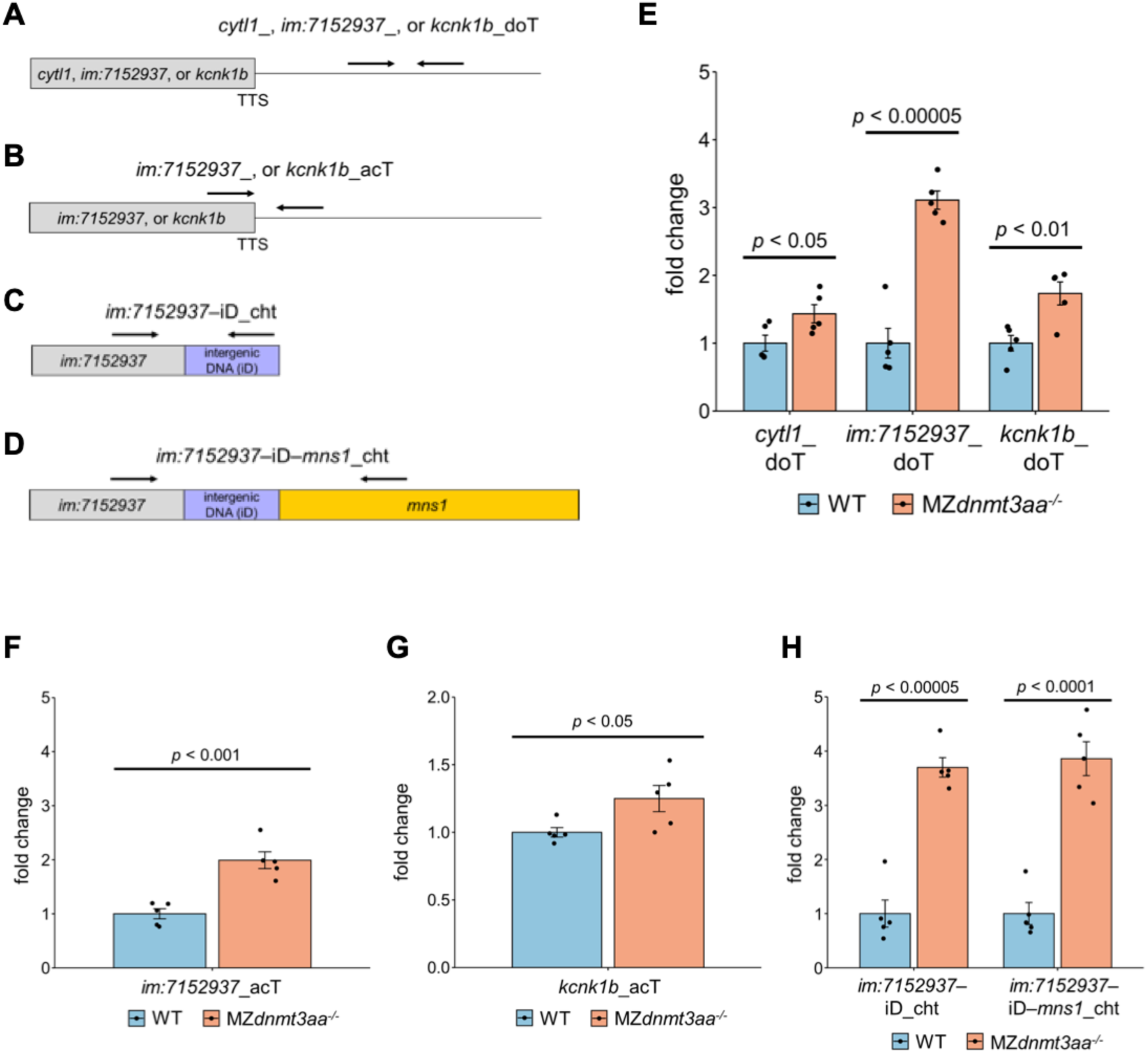
Identification of read-through and chimeric transcripts in MZ*dnmt3aa^-/-^* mutant. (A-D) Scheme of primer pair positions to detect aberrant transcripts. (E) qRT-PCR analysis for transcripts downstream of TTS (*cytl1*_doT, *im:7152937*_doT, and *kcnk1b*_ doT) in *cytl1*, *im:7152937*, and *kcnk1b* genes. (F, G) qRT-PCR analysis for transcripts across the TTSs (*im:7152937*_acT and *kcnk1b*_acT) in *im:7152937* (F) and *kcnk1b* (G) genes. (H) qRT-PCR analysis for chimeric transcripts between exons in *im:7152937* and iD/exon1 in *mns1* (*im:7152937*-iD_cht and *im:7152937*-*mns1*_cht). All qRT-PCR data are presented as means ± SE. The mean expression level of WT is defined as 1.

## Discussion

Unlike transcription initiation and elongation, the function of DNA methylation during transcription termination remains unclear. To investigate the DNA methylation function overlapping the TTS, we analyzed the impact of hypomethylation on transcription termination in *Dnmt3a*-deficient vertebrates. We showed that read counts downstream of TTS and number of chimeric transcripts with exons of downstream genes or iD had increased significantly in the *Dnmt3a*^-/-^ AgRPs, but not in *Dnmt3a^-/-^* ESCs and MEFs. Moreover, we identified increased read-through and chimeric transcripts in MZ*dnmt3aa^-/-^* zebrafish mutant. Taken together, we propose our model that DNA hypomethylation of TTSs is associated with the transcription termination defect in *Dnmt3a^-/-^* vertebrates (*Dnmt3a*^-/-^ AgRPs and MZ*dnmt3aa^-/-^* zebrafish mutant) (Figure 6).

**Figure 6.**
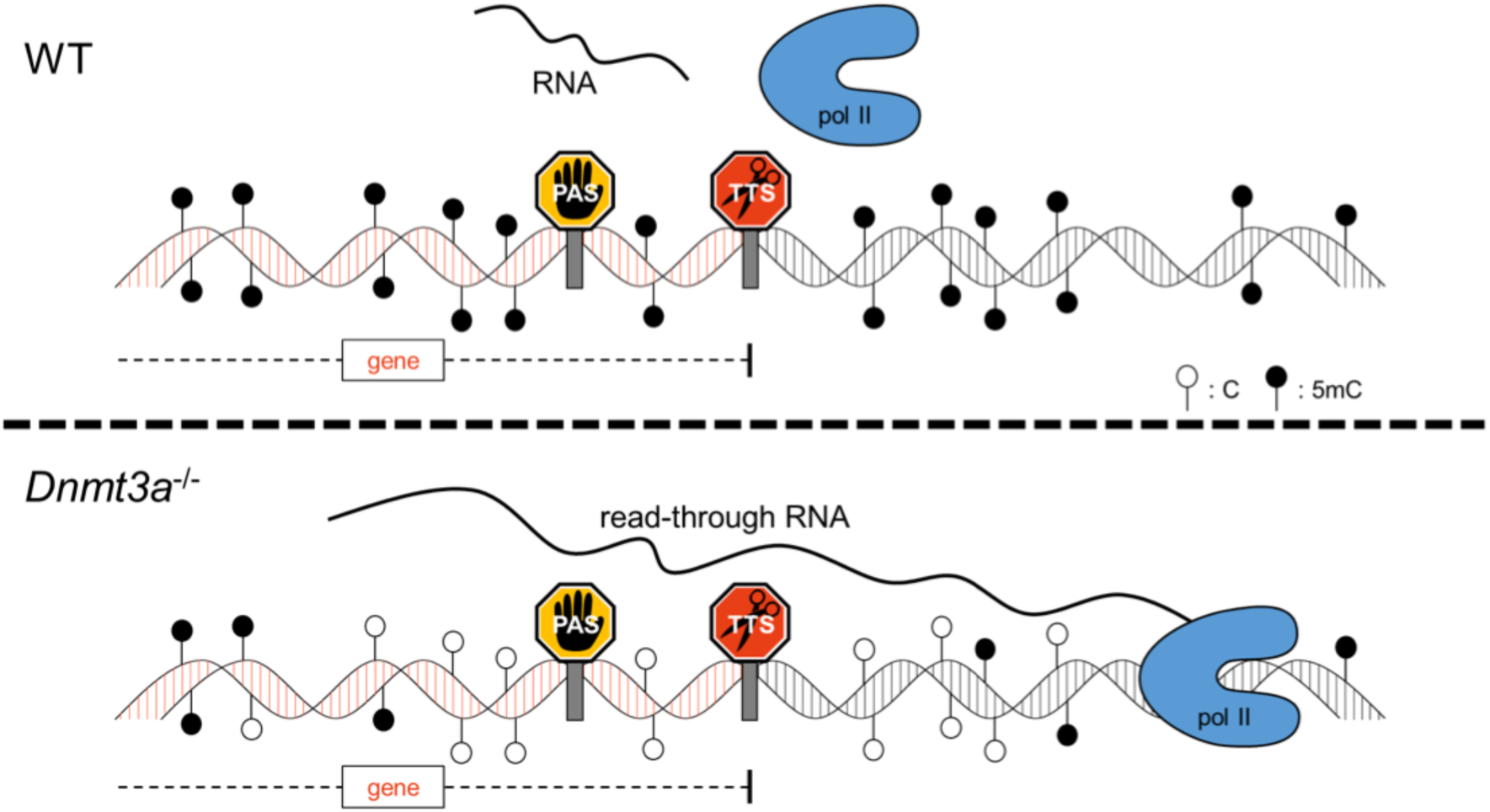
**A proposed model of transcription termination.**

Our bioinformatic analyses showed that the number of hypoDMRs overlapping TTSs in *Dnmt3a*^-/-^ J1 mESCs was much larger than that in *Dnmt3a*^-/-^ MEFs and AgRPs (Figure 1C). Previous studies have shown that mouse ESCs, such as J1 mESCs, which are cultured in medium containing serum, are globally hypermethylated compared to cells in mouse inner cell mass (Yagi *et al*. 2017b). In other words, *Dnmt3a* knockout J1 ESCs are thought to have a large number of hypoDMRs. The number of hypoDMRs overlapping TTSs in *Dnmt3a*^-/-^ J1 mESCs gradually decreased in the order of *Dnmt3a*^-/-^ J1 mESCs, MEFs, and AgRPs (Figure 1C). This result may be attributed to the fact that Dnmt3a regulates genome-wide methylation in ES cells; nevertheless, its role may be restricted as differentiation proceeds.

In this study, we used zebrafish mutant as an animal model because MZ*dnmt3aa^-/-^* mutant, unlike *Dnmt3a* knockout mice, are viable and fertile. The MZ*dnmt3aa^-/-^* mutant line can be easily maintained and in this mutant maternal Dnmt3aa functions can be excluded. However, normalized read counts downstream of the TTSs and chimeric transcripts were unchanged between WT and MZ*dnmt3aa^-/-^* mutant (Figure 4). There are two possible explanations for these results: one possibility is that zebrafish *dnmt3ab* may rescue the transcription termination defects in MZ*dnmt3aa^-/-^* mutant. Alternatively, it is hard to detect hypoDMRs overlapping TTSs in MZ*dnmt3aa^-/-^* mutant, because cells in the zebrafish body are more heterogeneous and vary greatly compared to AgRPs. Nevertheless, we have successfully detected the read-through and/or chimeric transcripts of three genes (*cytl1*, *im:7152937*, and *kcnk1b*) in MZ*dnmt3aa^-/-^* mutant, suggesting that DNA methylation is involved in transcription termination in zebrafish.

Although we have shown the significance of hypomethylation at the TTS region for transcription termination in vertebrates, it is unclear how hypomethylation regulates transcription termination. Despite the relative lack of transcription termination research, a unified allosteric/torpedo model has recently been proposed for the PAS-dependent termination (Eaton *et al*. 2020; Eaton and West 2020). In this model, the CPA complex recognizes and cleaves the nascent transcript when the pol II passes through the PAS; thus, PAS is close to TTS (10-35 bp upstream of the TTS) (Figure 6) (Tian and Manley 2017; Nourse *et al*. 2020). This cleavage by the CPA complex results in pol II pausing via dephosphorylation of the Spt5 elongation factor by a protein phosphatase 1 nuclear targeting subunit (PNUTS) (Eaton and West 2020). As paused pol II is dissociated from the DNA strand by the function of XRN2 exoribonuclease, which degrades RNA in a 5’→3’ direction, the pol II pausing is thought to be a critical step for the transcription termination (Proudfoot 2016; Eaton and West 2020). Although the CPA complex mainly binds to nascent pre-mRNA and C-terminal domain (CTD) of pol II, it is possible that DNA hypomethylation of the TTS or PAS genomic region may directly or indirectly suppress the CPA function, leading to loss of pol II pausing. It should be interesting to note that almost all hypoDMRs, which cause chimeric transcripts or increased read counts downstream of the TTSs, overlap with PAS in *Dnmt3a*^-/-^ AgRPs and MZ*dnmt3aa^-/-^* mutant (Figure 2H, Figure 4F, Supplementary Figure 5B, Supplementary Figure 8B). In PAS-dependent termination, there are two other mechanisms of pol II pausing: heterochromatin- and R-loop-dependent pausing (Proudfoot 2016). It has been reported previously that the heterochromatin structure downstream of the TTS causes pol II pausing by acting as an obstacle (Proudfoot 2016). Therefore, hypomethylation in the TTS region may lead to the suppression of heterochromatin formation downstream of TTS. An alternative possibility is R-loop-dependent pausing; R-loops are known as three-stranded structures, which consist of RNA-DNA hybrids, and function in various nuclear processes, such as transcription, DNA replication, and DNA repair (Al-hadid and Yang 2016; Niehrs and Luke 2020). As recent studies have shown that pol II pausing is induced by R-loop formation at the TTS, it is worth analyzing the relationship between R-loop formation and DNA methylation. Future molecular biological analyses regarding hypomethylation of the TTS region are needed to improve our understanding of the molecular events underlying transcription termination.

## Acknowledgements

We thank our laboratory members and Shyunya Hozumi for helpful discussion and critical comments; Masato Kinoshita for providing plasmids of CRISPR/Cas9 system; Mingcong Xu and Ikumi Taya for technical assistant. Computations were partially performed on the NIG supercomputer at ROIS National Institute of Genetics.

---

**Supplementary Figure S1.**
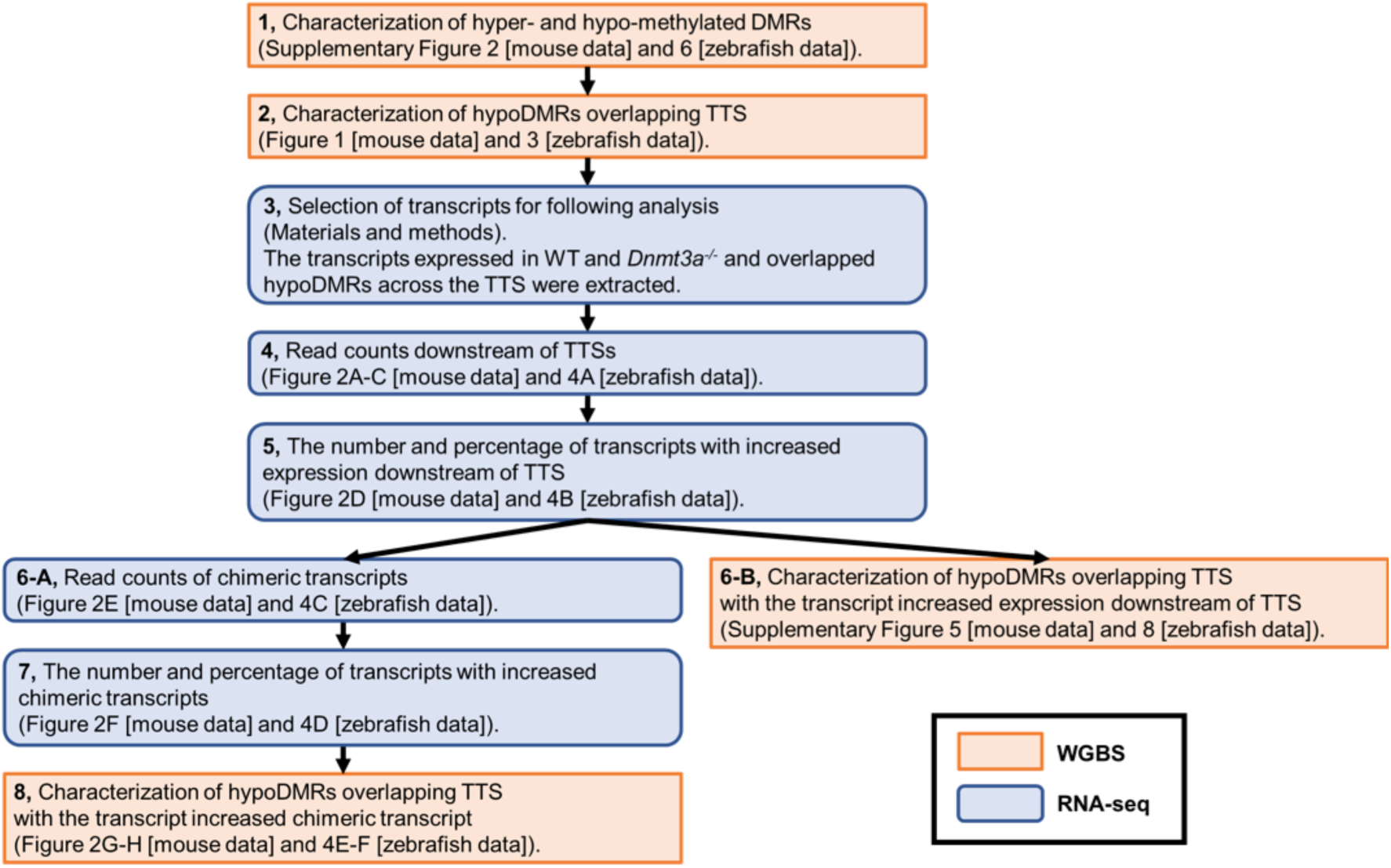
**Flow chart of bioinformatics analysis and experimental design.**

**Supplementary Figure S2.**
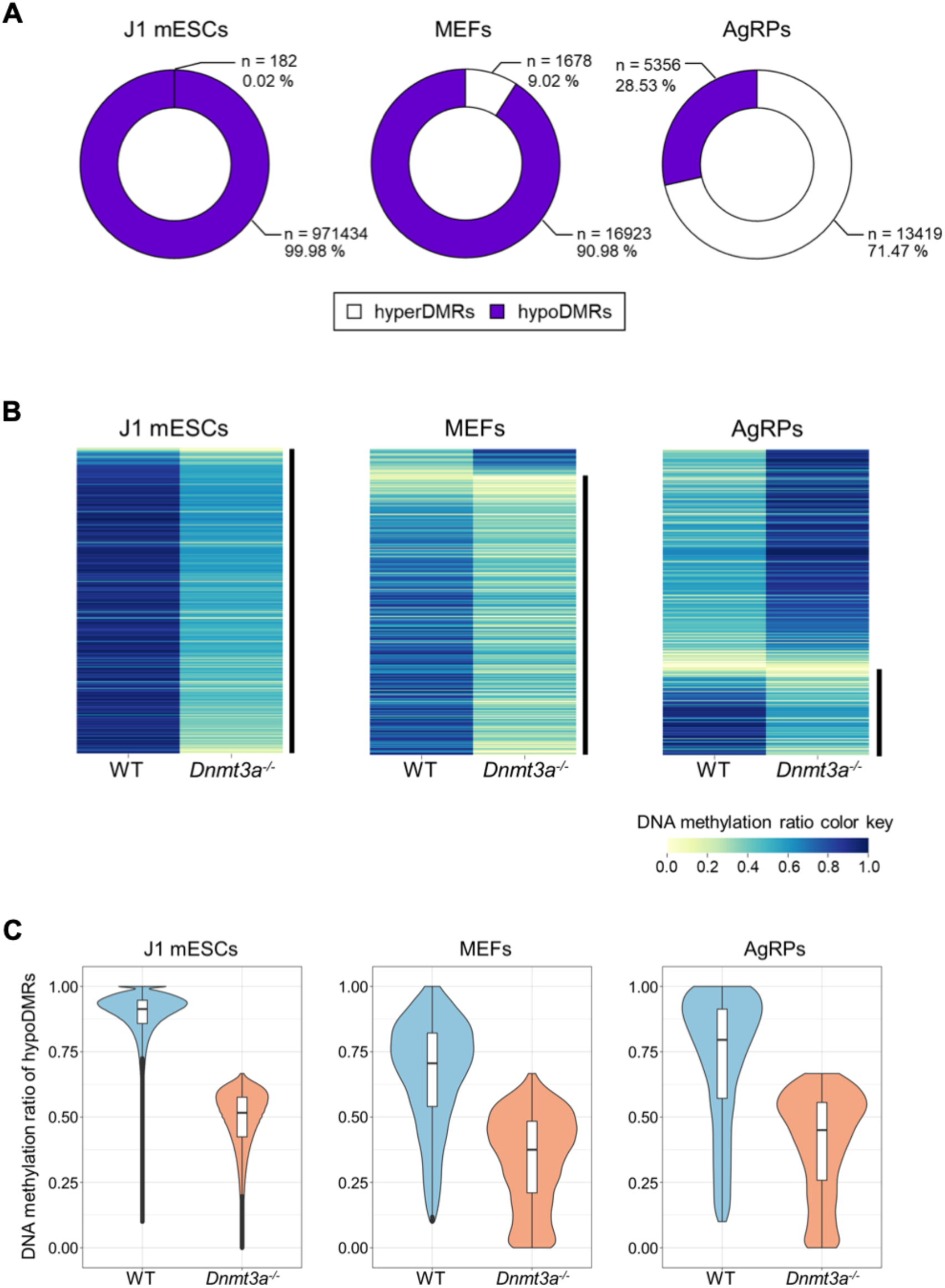
Characterization of hyperDMRs and hypoDMRs in three *Dnmt3a*-deficient mouse cells. (A) Pie charts depicting number and percentage of the hyperDMRs and hypoDMRs comparing WT with *Dnmt3a^-/-^* mutants (J1 mESCs, MEFs, and AgRPs). (B) Heatmaps of DNA methylation profiles of the DMRs comparing WT with *Dnmt3a^-/-^* mutants (J1 mESCs, MEFs, and AgRPs). Color scale indicates DNA methylation ratio. HypoDMRs are shown by vertical black bars. (C) Violin plots showing DNA methylation ratio of the hypoDMRs comparing WT with *Dnmt3a^-/-^* mutants (J1 mESCs, MEFs, and AgRPs).

**Supplementary Figure S3.**
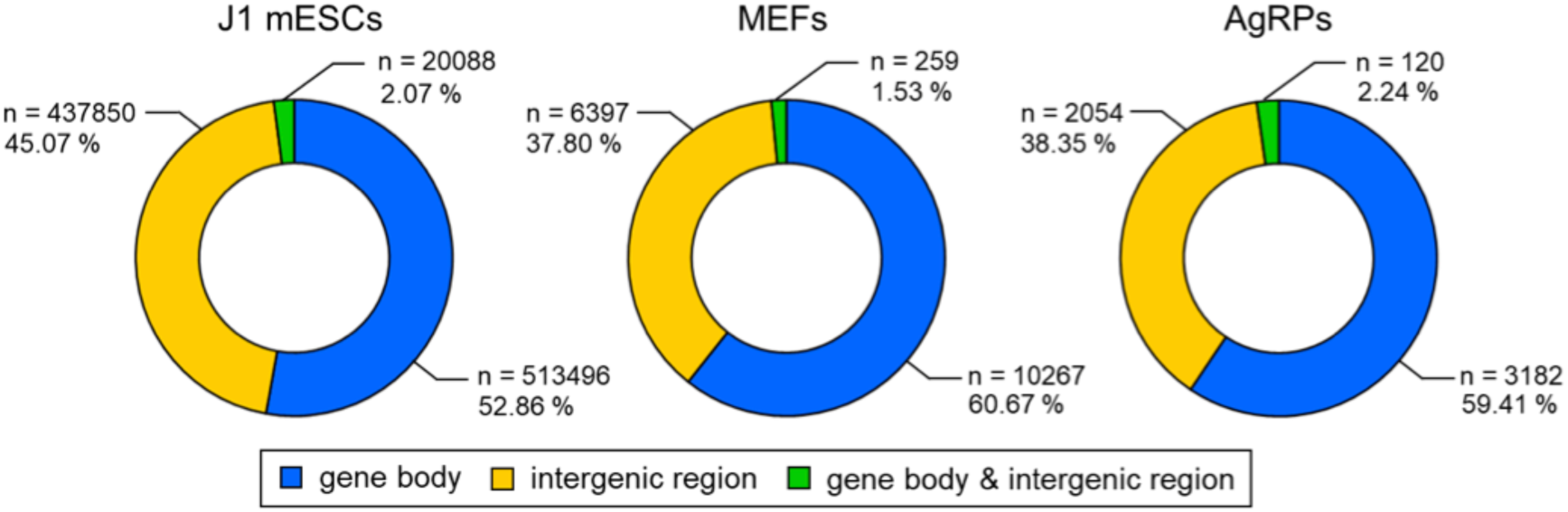
Genomic position of hypoDMRs in three *Dnmt3a* deficient mouse cells. Pie charts depicting number and percentage of genomic position (gene body, intergenic region, and gene body & intergenic region) of the hypoDMRs comparing WT with *Dnmt3a^-/-^* mutants (J1 mESCs, MEFs, and AgRP).

**Supplementary Figure S4.**
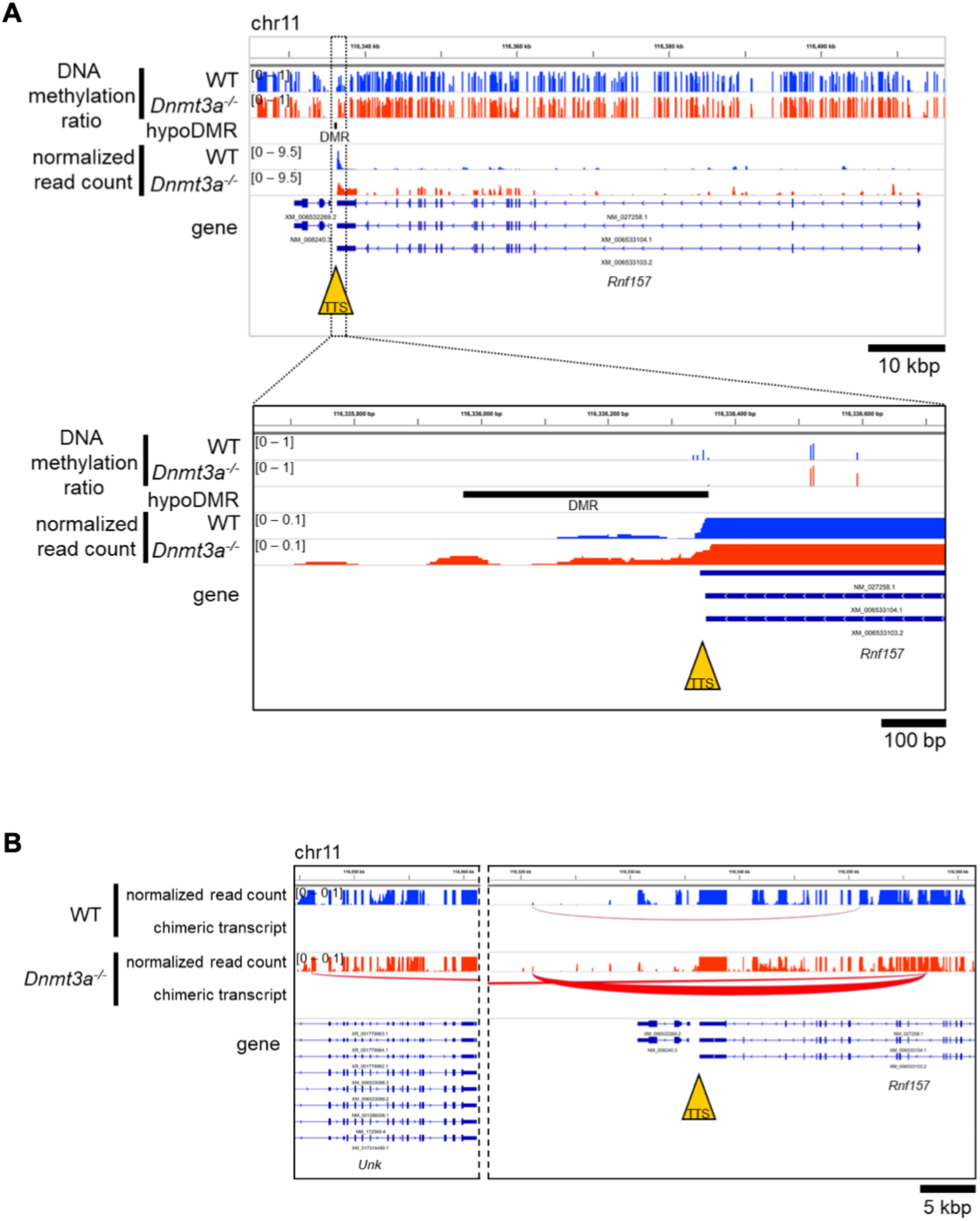
Transcription termination defects of *Rnf157* gene in *Dnmt3a^-/-^* AgRPs. (A, B) Genome browser view for *Rnf157* gene in *Dnmt3a^-/-^* AgRPs. Normalized read counts (A) and chimeric transcripts (B), as an example, are shown in this Figure.

**Supplementary Figure S5.**
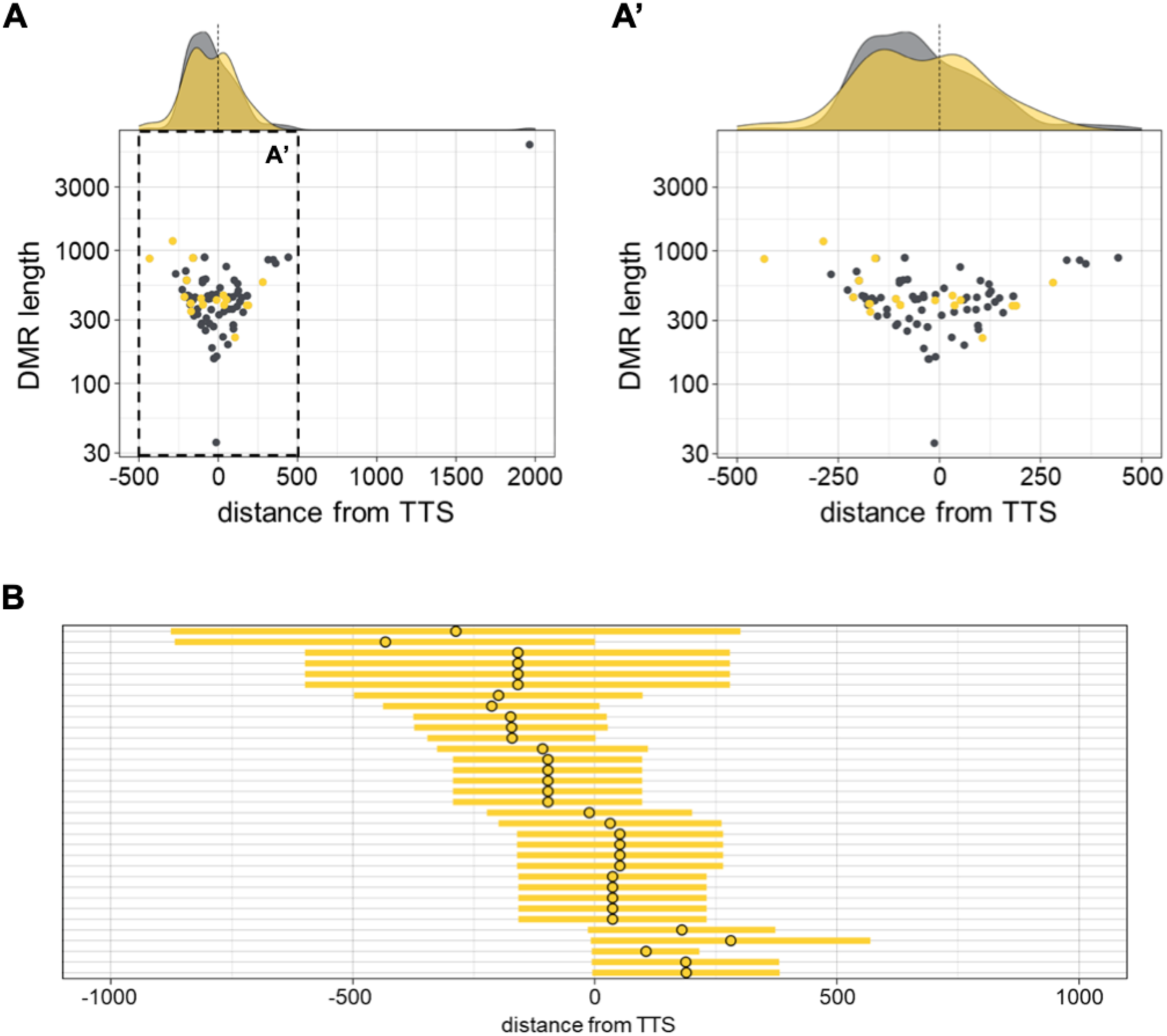
Distribution of hypoDMRs which lead to increased read counts downstream of TTSs in *Dnmt3a^-/-^* AgRPs. (A, A’) 2D scatterplots with marginal distribution comparing hypoDMR length versus distance from TTS in *Dnmt3a^-/-^* AgRPs. Grey and yellow dots represent the center of hypoDMRs. Yellow dots represent the hypoDMRs which lead to increased read counts downstream of TTSs. A right graph (A’) is an enlarged view of the left one (A). (B) A horizontal bar graph representing the distribution of hypoDMRs, which lead to increased read counts, relative to TTS. Yellow dots and bars represent the center and range of hypoDMRs, respectively.

**Supplementary Figure S6.**
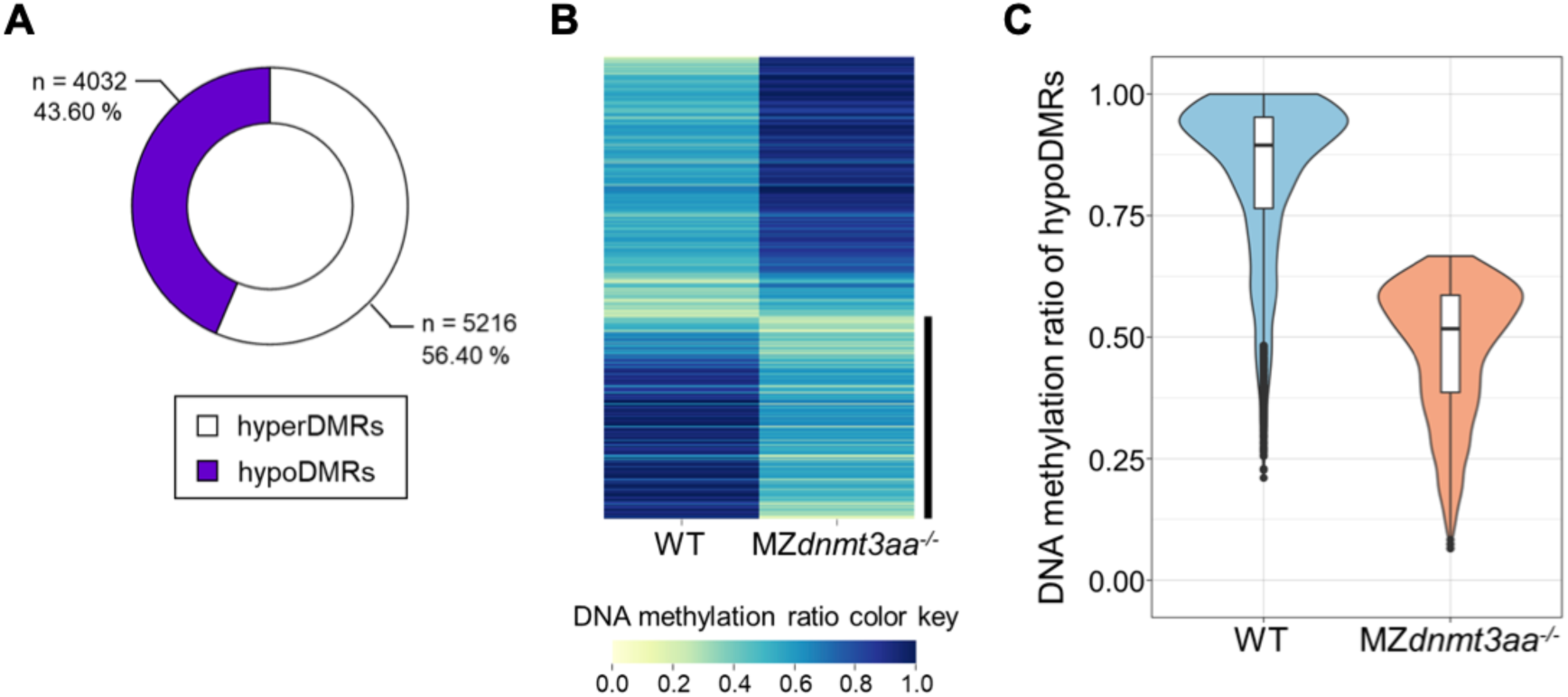
Characterization of hyperDMRs and hypoDMRs in MZ*dnmt3aa^-/-^* mutant. (A) A pie chart depicting number and percentage of the hyperDMRs and hypoDMRs comparing WT with MZ*dnmt3aa^-/-^* mutant. (B) A heatmap of DNA methylation profiles of the DMRs comparing WT with MZ*dnmt3aa^-/-^* mutant. (C) A violin plot showing DNA methylation ratio of the hypomethylated DMRs comparing WT with MZ*dnmt3aa^-/-^* mutant.

**Supplementary Figure S7.**
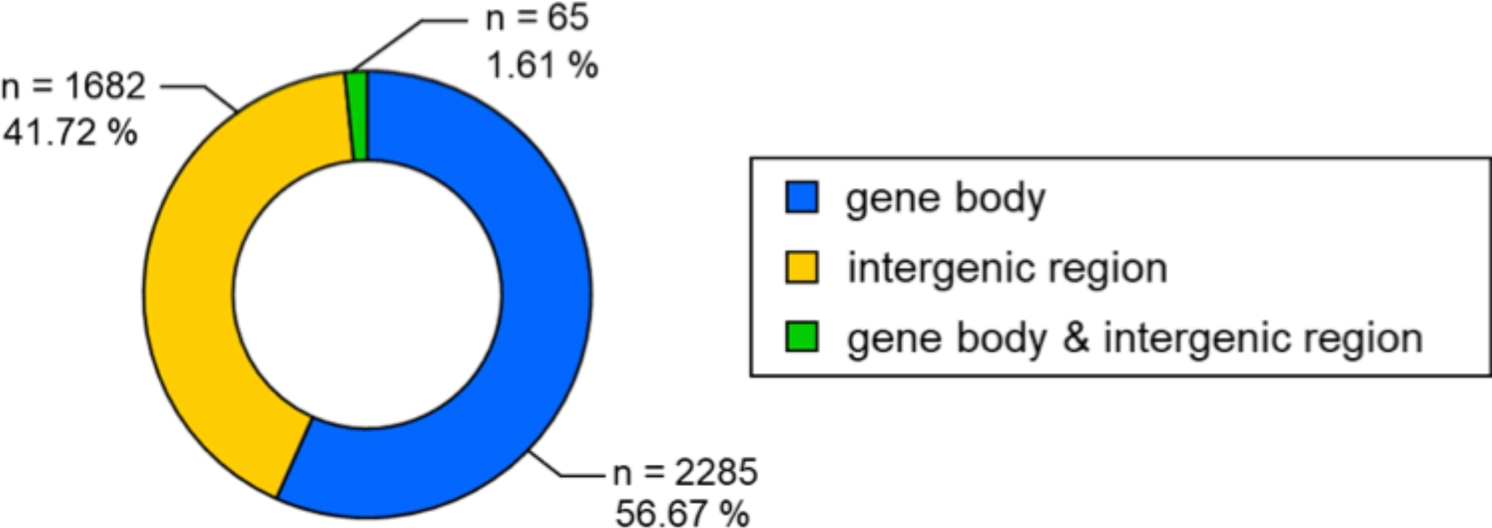
Genomic position of hypoDMRs in MZ*dnmt3aa^-/-^* mutant. A pie chart depicting number and percentage of genomic position (gene body, intergenic region, and gene body & intergenic region) of the hypoDMRs comparing WT with MZ*dnmt3aa^-/-^* mutant.

**Supplementary Figure S8.**
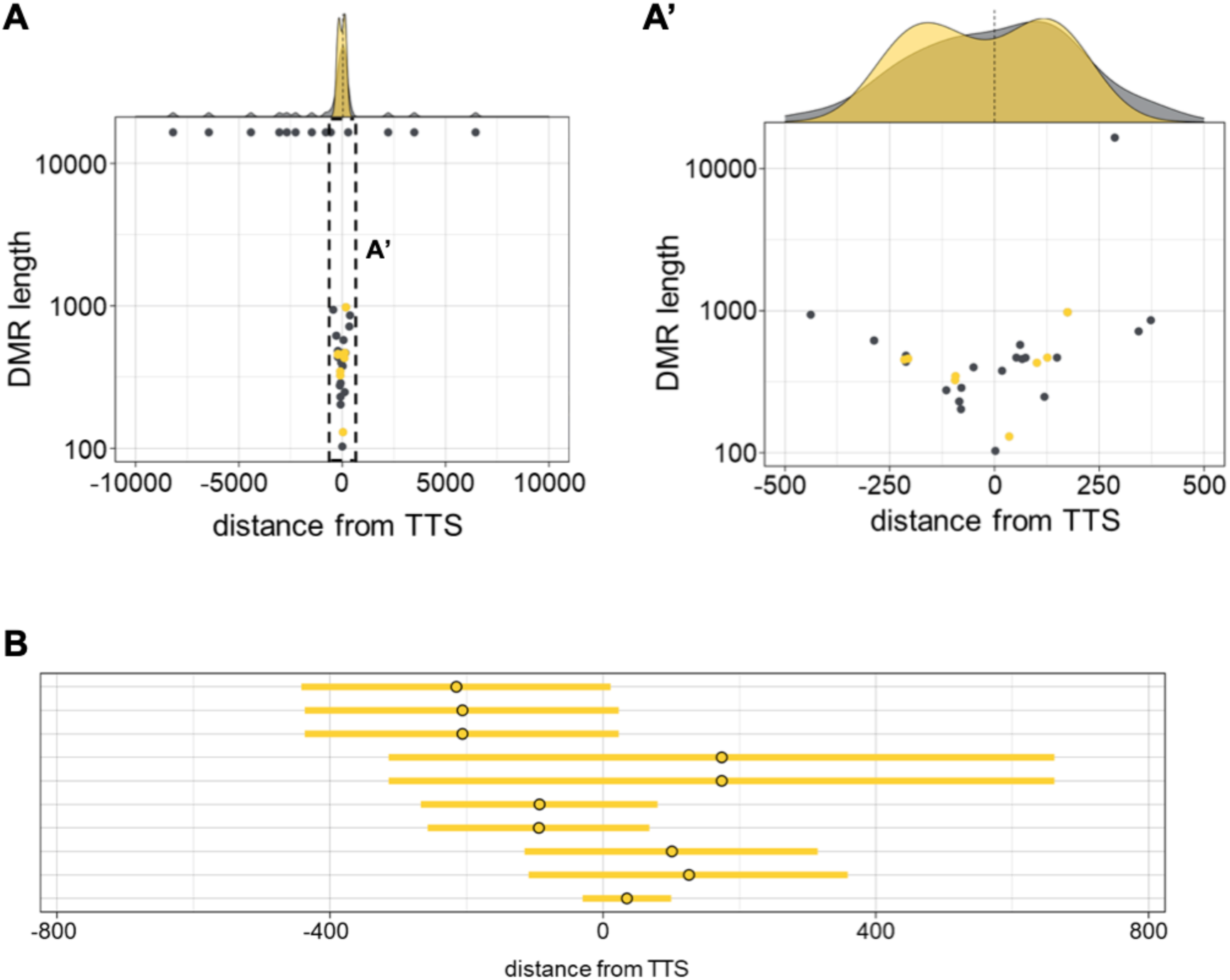
Distribution of hypoDMRs which lead to increased read counts downstream of TTSs in MZ*dnmt3aa^-/-^* mutant. (A, A’) 2D scatterplots with marginal distribution comparing hypoDMR length versus distance from TTS in MZ*dnmt3aa^-/-^* mutant. Grey and yellow dots represent the center of hypoDMRs. Yellow dots represent hypoDMRs which lead to increased read counts downstream of TTSs. A right graph (A’) is an enlarged view of the left one (A). (B) A horizontal bar graph represents the distribution of hypoDMRs, which lead to increased read counts, relative to TTS. Yellow dots and bars represent the center and range of hypoDMRs, respectively.

**Supplementary Figure S9.**
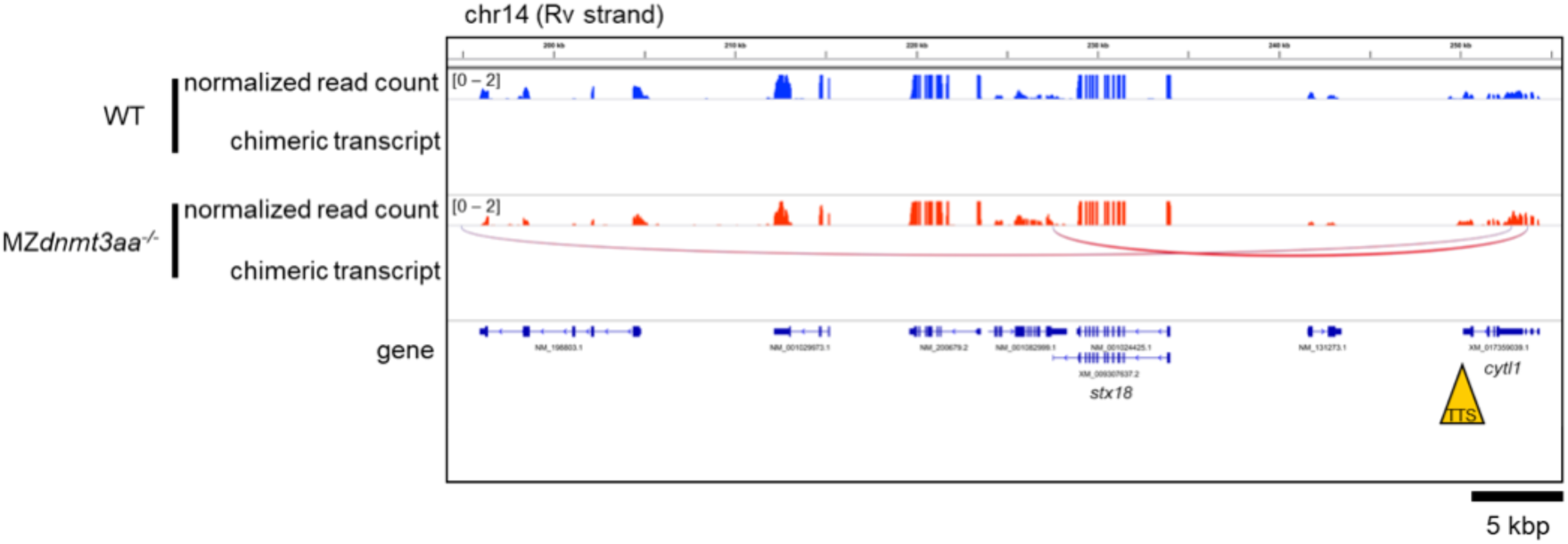
Transcription termination defects of *cytl1* gene in MZ*dnmt3aa^-/-^* mutant. Genome browser view for *cytl1* gene in MZ*dnmt3aa^-/-^* mutant. Chimeric transcripts, as an example, are shown in this Figure.

**Supplementary Figure S10.**
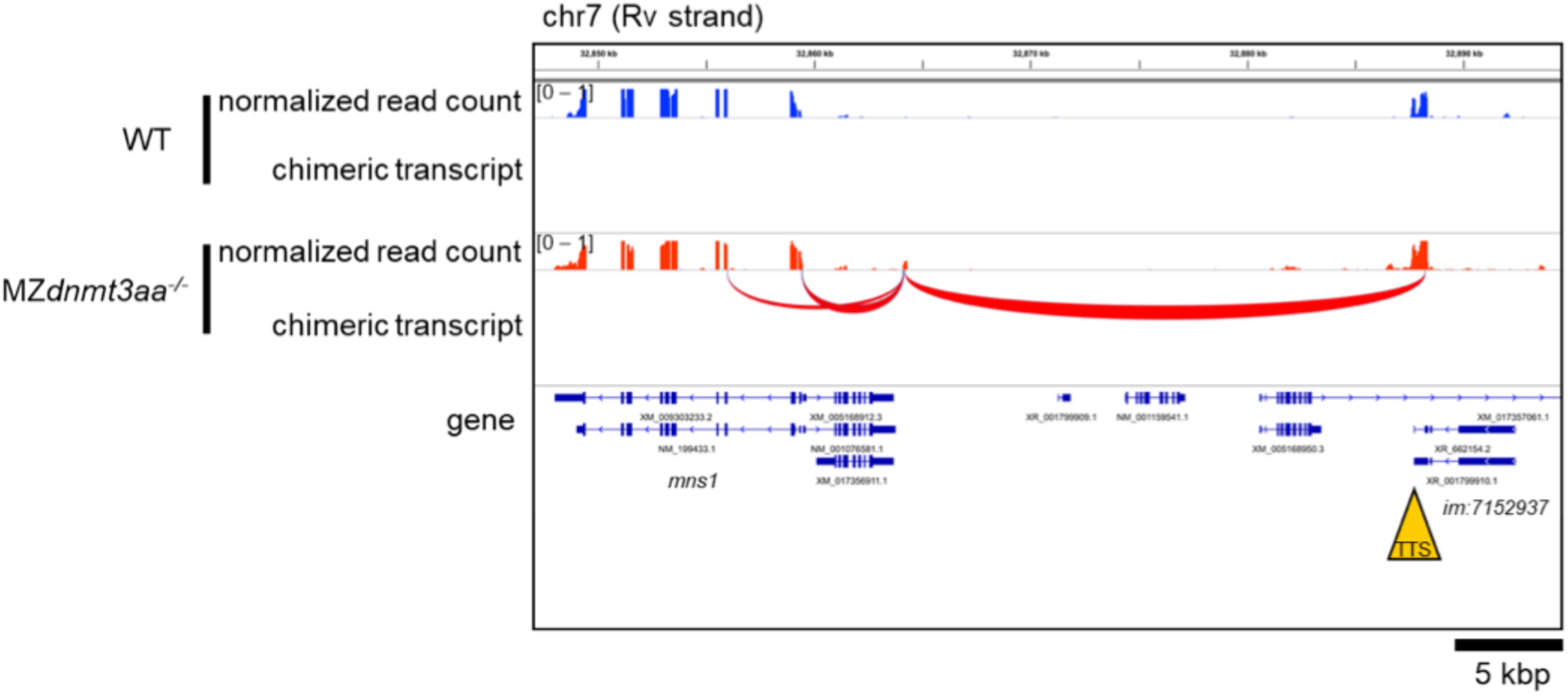
Transcription termination defects *im:7152937* in MZ*dnmt3aa^-/-^* mutant. Genome browser view for *im:7152937* in MZ*dnmt3aa^-/-^* mutant. Chimeric transcripts, as an example, are shown in this Figure.

**Figure S11.**
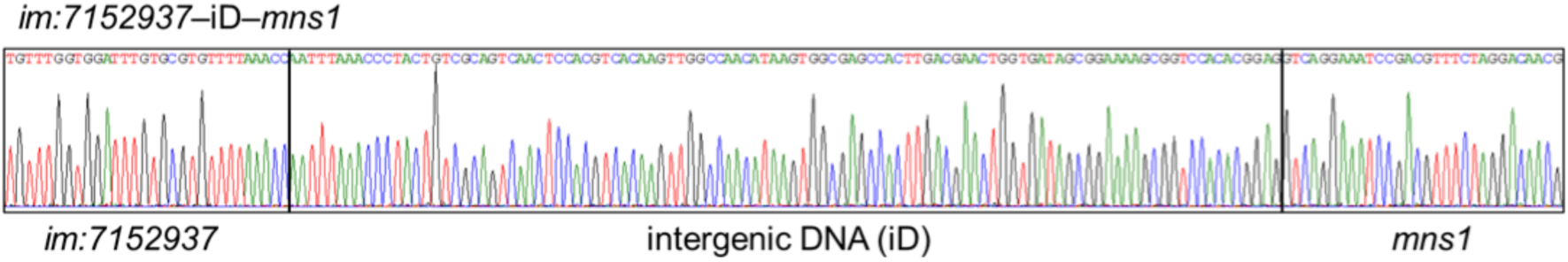
A sequencing chromatogram of chimeric transcripts, *im:7152937*-iD-*mns1*.

**Supplementary Table S1.**
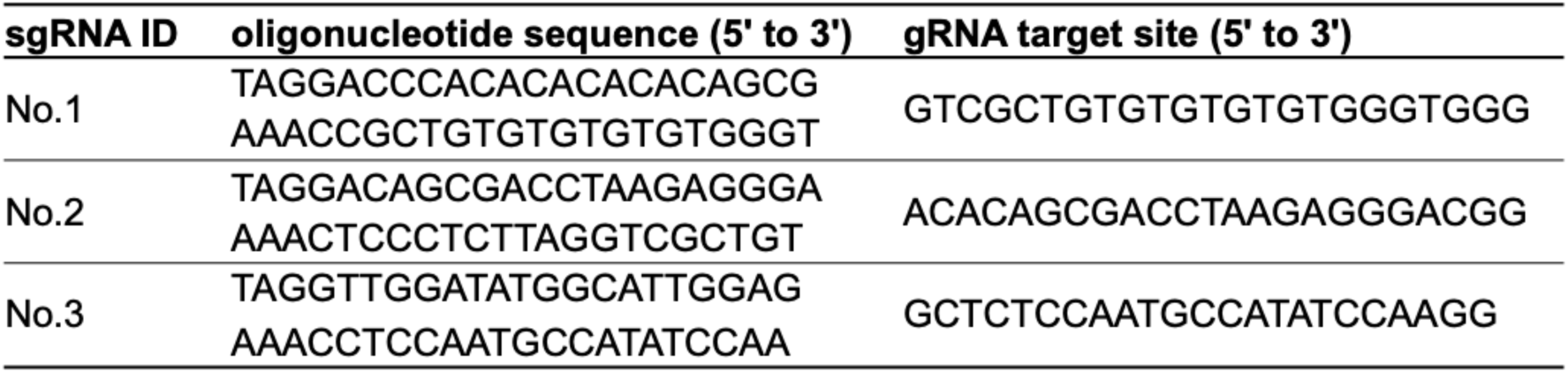

**Supplementary Table S2.**
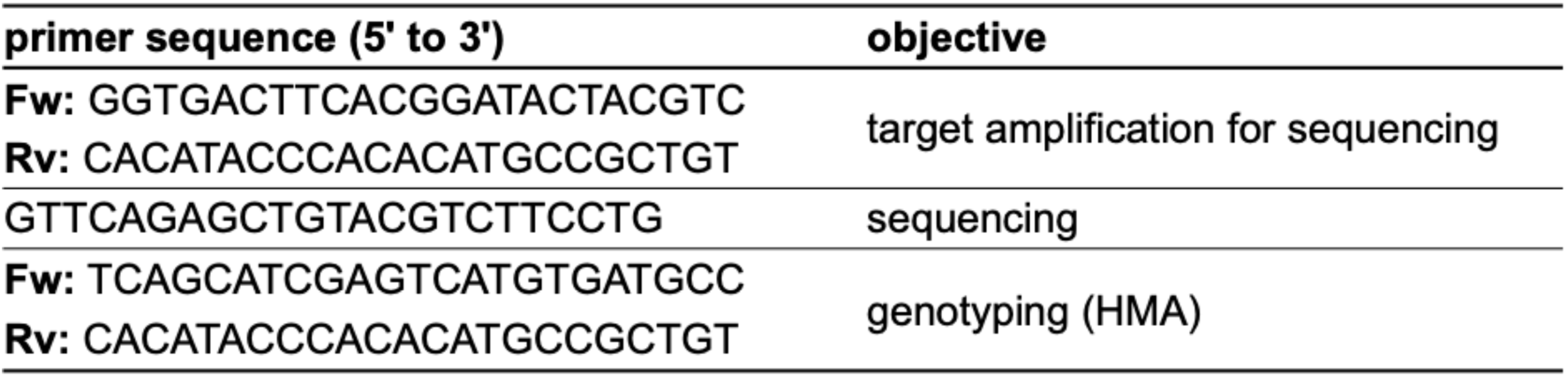

**Supplementary Table S3.**
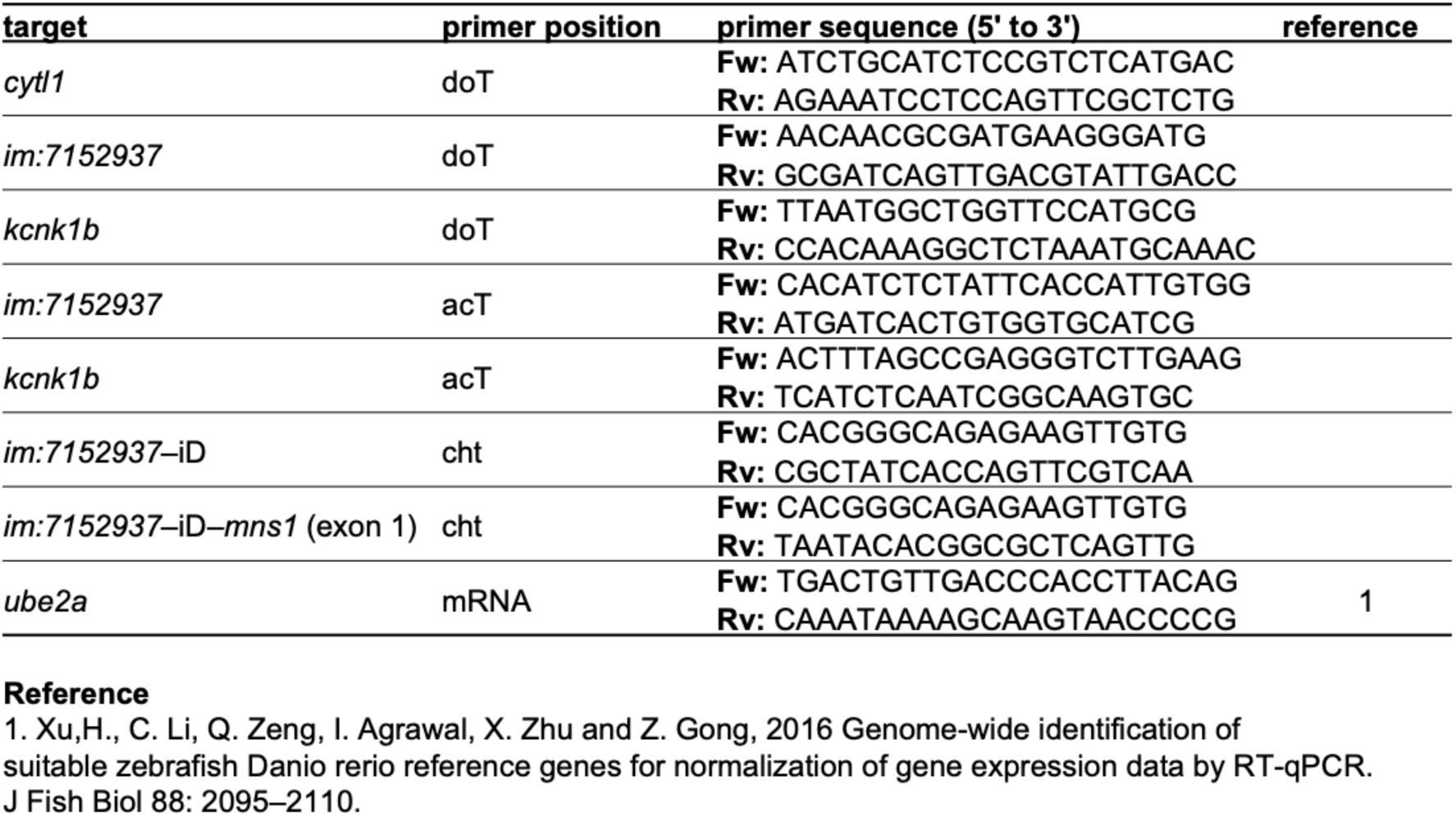

**Supplementary Table S4.**
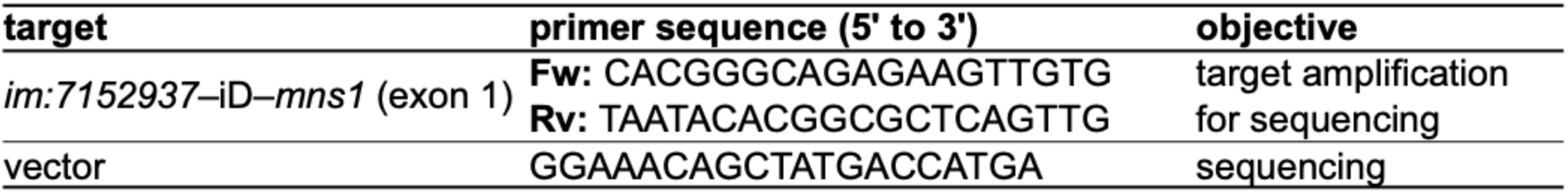

